# Post-mitotic transcriptional activation and 3D regulatory interactions show locus- and differentiation-specific sensitivity to cohesin depletion

**DOI:** 10.1101/2025.02.13.638153

**Authors:** UkJin Lee, Alejandra Laguillo-Diego, Wilfred Wong, Zhangli Ni, Lingling Cheng, Jieru Li, Bobbie Pelham-Webb, Alexandros Pertsinidis, Christina Leslie, Effie Apostolou

**Affiliations:** Sanford I. Weill Department of Medicine, Sandra and Edward Meyer Cancer Center, Weill Cornell Medicine, New York, NY 10021, USA; Molecular Biology Program, Graduate School of Medical Sciences, Weill Cornell Medicine, New York, 10065, USA; Computational and Systems Biology Program, Memorial Sloan Kettering Cancer Center, New York, NY 10065, USA; Tri-Institutional Training Program in Computational Biology and Medicine, New York, NY 10065, USA; Structural Biology Program, Memorial Sloan Kettering Cancer Center, New York, NY 10065, USA; Weill Cornell/Rockefeller/Sloan Kettering Tri-Institutional MD-PhD Program, New York, NY 10021, USA

## Abstract

Prior studies showed that structural loops collapse upon acute cohesin depletion, while regulatory enhancer-promoter (E-P) loops largely persist, consistent with minimal transcriptional changes. However, these studies, conducted in asynchronous cells, could not resolve whether cohesin is required for the establishment of regulatory interactions and transcriptional activation during cell division or cell state transitions. To address this gap, we degraded RAD21, a core cohesin subunit, in naïve mouse embryonic stem cells (ESCs) transitioning from mitosis to G1 either in self-renewal condition or during differentiation toward formative pluripotency. Although most structural loops failed to be re-established without cohesin, about 35% of regulatory loops reformed at normal or higher frequencies. Cohesin-independent loops showed characteristics of strong active enhancers and promoters and a significant association with H3K27ac mitotic bookmarks. However, inhibition of CBP/p300 during mitotic exit did not impact these cohesin-independent interactions, suggesting the presence of complex compensatory mechanisms. At the transcriptional level, cohesin depletion induced only minor changes, supporting that post-mitotic transcriptional reactivation is largely independent of cohesin. The few genes with impaired reactivation were directly bound by RAD21 at their promoters, engaged in many structural loops, and located within strongly insulated TADs with low gene density. Importantly, degrading cohesin during the M-to-G1 transition in the presence of EpiLC differentiation signals revealed a larger group of susceptible genes, including key signature genes and transcription factors. Impaired activation of these genes was partly due to the failure to establish *de novo* EpiLC-specific interactions in the absence of cohesin. These experiments revealed locus-specific and context-specific dependencies between cohesin, E-P interactions, and transcription.

## Introduction

Cell fate regulation relies on the orchestrated activation of enhancers and their target genes by cell-type specific transcription factors in the context of the three-dimensional (3D) nucleus.^1^ 3D chromatin conformation technologies have revealed that the genome is hierarchically organized into compartments, topologically associated domains (TADs), and chromatin loops, which bring distal genomic elements, such as enhancers and promoters, into spatial proximity.^2,3^ These structures are highly dynamic and undergo extensive reorganization throughout the cell cycle^4–10^, during development^11–16^, and in disease.^17–19^ They are tightly linked to cell-type-specific transcriptional programs, with evidence suggesting they can restrict, support, or instruct enhancer-promoter (E-P) communication and gene expression.^20–22^ Despite these strong associations between 3D chromatin folding and transcription, the functional interdependence between these processes and their driving forces remain highly debated.^23–25^

One central regulator of the 3D chromatin architecture is cohesin, a ring-shaped protein complex with an ATP-dependent “motor-like” activity that enables DNA extrusion into loop-like structures.^26–29^ Cohesin aberrations can lead to disorders known as cohesinopathies^18,19^ but also contribute to a variety of cancers.^17^ Before the discovery of cohesin’s role in organizing interphase 3D chromatin organization, cohesin was mostly known for its essential role in sister-chromatid cohesion during mitosis (M).^30^ This dual role presented a challenge for the proper interpretation of results and phenotypes arising from long-term perturbations of cohesin. To overcome these limitations, recent studies in various cellular contexts employed acute protein degradation approaches for various cohesin components, including the core subunit RAD21, for durations shorter than a cell cycle.^31–39^ These studies revealed dramatic effects on the 3D chromatin organization with a nearly complete abrogation of TADs and loops and a reciprocal strengthening of compartmentalization. Intriguingly, despite these major architectural perturbations, the impact on global transcriptional activity was limited, suggesting that cohesin-mediated chromatin organization and transcription are largely uncoupled. More recent experiments employing higher-resolution 3D genomics approaches, which more effectively capture E-P loops, have documented that a large number of regulatory interactions can persist—at least for short periods of time—in the absence of cohesin, suggesting the presence of compensatory, cohesin-independent mechanisms.^20,33,39^ Although these results strongly support that cohesin is largely dispensable for the maintenance of E-P interactions and transcriptional activity, whether this is true for *de novo* establishment of chromatin loops and transcriptional activation remains unknown.

Every time a cell goes through mitosis (M), its 3D chromatin organization and transcriptional activity are drastically perturbed, along with the dissociation of many chromatin factors—including extruding cohesin—from the mitotic genome.^4–7,40–42^ Upon mitotic exit and during G1 transition (M-to- G1), these cell-type defining features need to be rapidly and faithfully reset for proper inheritance/self-renewal of cell identity.^8,9,40,41,43^ Therefore, the M-to-G1 transition constitutes a window of vulnerability for cell fate change^44–46^, especially for progenitors or embryonic stem cells (ESCs) that constantly balance self-renewal and differentiation. Moreover, the M-to-G1 transition also offers a unique system to dissect the interplay between 3D organization and transcription. Previous studies from our group and others have characterized, at a genome-wide scale, the dynamic transcriptional and architectural resetting that occurs during the M-to-G1.^6–8,10^ In mouse ESCs, transcription is reactivated in waves with a rapid re-induction of stem cell-associated genes and enhancers, followed by the transient activation/de-repression of lineage-specific genes, suggesting transient priming towards differentiation.^8^ Topological reorganization at different hierarchical levels (compartments, TADs, and loops) also happens asynchronously. A/B compartmentalization appears early after mitotic exit and becomes progressively stronger throughout G1.^6–8^ Similarly, TAD boundaries emerge early, initially defining smaller TADs that merge into larger, nested TADs.^6–8^ At a finer scale, regulatory interactions reform faster than structural interactions between CTCF/cohesin anchors.^7,8^ To what degree is cohesin required for the post-mitotic establishment of any of these architectural and transcriptional features remains an open question.

Here, we exploit the unique M-to-G1 “resetting” window in synchronized mouse ESCs in combination with acute perturbations to determine the cohesin-dependent and independent aspects of the post-mitotic 3D organization and transcriptional reactivation. Importantly, we perform these experiments both in self-renewing ESCs (naïve pluripotency) and acute differentiation conditions (towards epiblast-like cells (EpiLCs) representing formative pluripotency). This approach allowed us to assess the relative impact of cohesin depletion not only on the post-mitotic reactivation of the cell-type-specific transcriptional programs but also on the establishment of novel transcriptional programs and 3D chromatin topologies associated with new cell identity. Our findings support a critical role of cohesin on the post-mitotic 3D chromatin re-organization but a limited and locus-specific impact on the re-formation of E-P interactions and transcriptional activation. Our results were concordant with previous studies in asynchronous cells^33^ with no evidence for a preferential vulnerability during establishment (M-to-G1) rather than maintenance (interphase). Importantly, RAD21 depletion during the M-to-G1 transition in the presence of differentiation signals revealed more extensive and pronounced effects for newly activated genes and *de novo* established E-P contacts, leading to impaired cell fate transitions. Together, our results support that both the establishment and maintenance of transcriptional activity are largely independent of cohesin. However, we also uncover context- and locus-specific vulnerabilities and describe the key features and principles of cohesin-dependent or independent chromatin interactions and transcription, as well as potential compensatory mechanisms.

## Results

### A system for acute cohesin depletion without interfering with the M-to-G1 transition

To study the role of cohesin-mediated loop extrusion in the resetting of 3D chromatin architecture and transcription during the M-to-G1 phase, we used mouse embryonic stem cells (ESCs; *Rad21*- dTAG cell line) engineered with the dTAG system^47^ for inducible degradation of RAD21, a core subunit of the cohesin complex (**Fig. 1a**). RAD21 was homozygously tagged with the FKBP12^F36V^ degron, enabling its acute depletion upon dTAG-13 treatment (hereafter referred to as dTAG)^48^. The RAD21 construct also included two HA tags and a HaloTag, facilitating downstream applications such as Western blot and imaging. Treatment with dTAG for various durations revealed ∼90% degradation efficiency at 2 hours and complete depletion after that timepoint as quantified by Western blot experiments (**Extended Data** Fig. 1a). Importantly, dTAG-induced RAD21 degradation was evident both in the nuclear soluble and chromatin-bound fractions, supporting that our system effectively depletes chromatin-engaged, loop-extruding cohesin (**Extended Data** Fig. 1b).

**Fig. 1.**
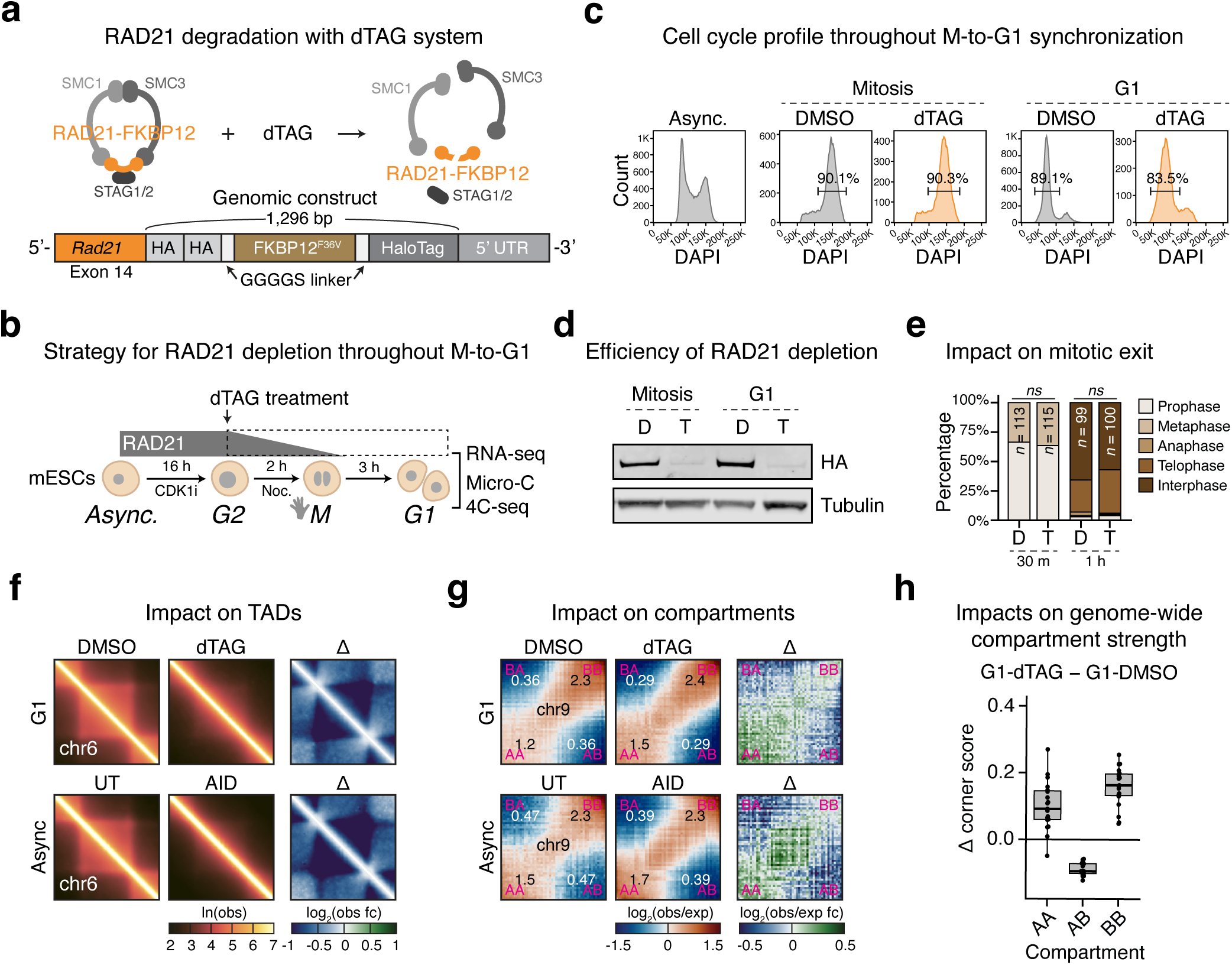
| Acute cohesin depletion during the M-to-G1 phase in mouse ESCs recapitulates higher- order 3D chromatin architecture changes seen in asynchronous perturbation. a, Top panel: Schematic of the RAD21 depletion strategy, in which dTAG-13 (dTAG) treatment induces the degradation of the RAD21-FKBP12 fusion protein, leading to the disruption of the cohesin complex. Bottom panel: Illustration of the knock-in cassette introduced into the endogenous *Rad21* gene locus in mouse ESCs, positioned after the last exon, enabling inducible RAD21 depletion via the dTAG system. The RAD21 protein is fused to the FKBP12^F36V^ peptide, flanked by GGGGS linkers, along with two HA tags and a HaloTag. b, Experimental strategy for dTAG-induced RAD21 depletion during the M-to-G1 transition. ESCs were synchronized in mitosis by treatment with 9 µM CDK1 inhibitor for 16 hours, followed by 100 ng/mL nocodazole treatment for 2 hours. After mitotic shake-off, cells were released into G1 for 3 hours. To ensure RAD21 depletion upon G1 entry, cells were treated with 500 nM dTAG starting at G2, with 0.05% v/v DMSO used as a control. Cells in G1 were collected for RNA- seq, Micro-C, and 4C-seq. c, Representative cell cycle profile analyzed by FACS using DAPI staining, highlighting a prominent 4N peak (with percentage) in mitosis and a prominent 2N peak (with percentage) in G1, comparing dTAG-treated ESCs with the DMSO control. d, Representative Western blot images showing near-complete degradation of RAD21 (anti-HA) in ESCs in both mitosis and G1 following dTAG treatment. DMSO (D). dTAG (T) e, Stacked bar plot showing the percentage of ESCs in different cell cycle phases at 30 min and 1 hour after mitotic release, following continuous 0.05% v/v DMSO (D) or 500 nM dTAG (T) treatment from G2. Cells were assigned to different cell cycle phases based on immunofluorescence analysis using DAPI and antibodies against H3S10ph and tubulin (Extended Data Fig. 1e). The number of cells counted per condition is indicated. No significant differences were observed between DMSO and dTAG conditions (Fisher’s exact test, *ns* = not significant). Representative data from two independent experiments are shown. f–h, Architectural changes based on Micro-C data analysis from G1 (this study) and asynchronous (async)^33^ ESCs with or without RAD21 depletion. f, Left panel: Representative aggregate TAD plots from chr6 in DMSO- and dTAG-treated G1 cells, as well as UT (control) and AID (RAD21-depleted) asynchronous conditions. Color represents natural log (ln) of the observed (obs) contact frequency. Right panel: Differential (Δ = dTAG − DMSO) plot comparing RAD21-depleted and control conditions. Color represents log_2_ fold change in observed (obs) contact frequency. g, Left panel: Representative saddle plots from chr9 in DMSO- and dTAG-treated G1 cells, as well as UT (control) and AID (RAD21- depleted) asynchronous conditions. The average signal intensities for the 10% area at each corner are displayed. Color represents log_2_ of observed (obs) contact frequency over expected (exp) contact frequency. Right panel: Differential (Δ = dTAG − DMSO) plot comparing RAD21-depleted and control conditions. Color represents log_2_ fold change in observed (obs) contact frequency over expected (exp) contact frequency. h, Box plot showing the intra- (AA or BB) or inter (AB) compartmental changes across all chromosomes upon RAD21 depletion. The y-axis shows the difference (Δ = dTAG − DMSO) of the saddle plot corner scores (as shown in panel g). Boxes represent the interquartile range, with lines indicating the median. Each data point corresponds to an individual chromosome.

To specifically deplete cohesin during the M-to-G1 transition, we first synchronized ESCs in the G2 phase with a CDK1 inhibitor (CDK1i) and then treated the cells with both nocodazole and dTAG (or DMSO as control) for 2 hours to induce mitotic arrest and RAD21 degradation (**Fig. 1b**). After mitotic shake-off and extensive washout of the nocodazole, cells were released in G1 for 3 hours in the presence of dTAG to ensure continuous cohesin depletion before collection for transcriptomics and genomics assay. The efficiency of mitotic arrest and release was quantified by FACS for DNA content (DAPI) (**Fig. 1c**) and by immunofluorescence (IF) staining for the mitotic mark H3S10ph (**Extended Data** Fig. 1c), and only samples with >90% mitotic purity and >80% mitotic release into G1 were used for downstream experiments. Importantly, we also confirmed by Western blot that RAD21 was fully degraded in the collected G1 samples after dTAG treatment (**Fig. 1d**).

A major concern when depleting RAD21 is that cohesin is required for sister chromatid cohesion during mitosis, and its loss could disrupt proper mitotic progression. However, we observed very similar efficiencies of mitotic arrest and release into G1 both in the DMSO and dTAG conditions (**Fig. 1c** and **Extended Data** Fig. 1c) despite the complete depletion of RAD21 at the time of the collection (**Fig. 1d**). Previous studies in other organisms have shown that sister chromatid cohesion and chromatin segregation occur normally with up to 80–90% cohesin depletion^49,50^. Therefore, we wondered to what degree the remaining ∼10% of RAD21 at the time of the mitotic shake-off (upon 2 hours of dTAG treatment, as shown in **Extended Data** Fig. 1a) might be sufficient to support normal mitotic exit prior to complete RAD21 depletion. To test this, we first analyzed the impact of dTAG treatment on the cell cycle profile of asynchronous mESCs (**Extended Data** Fig. 1d). We observed that cells started to gradually accumulate in the G2/M phase (4N DNA content) after 2 hours of treatment, indicating minimal impact on mitotic exit up to this point. To further assess the impact of RAD21 depletion on mitosis, we collected mitotically arrested ESCs with or without dTAG treatment and quantified their progress toward G1, 30 minutes and 1 hour after extensively washing out nocodazole and dTAG. Immunofluorescence analysis documented no significant differences (Fisher’s exact test) between DMSO and dTAG-treated samples in any of the mitotic stages (prophase, metaphase, anaphase, and telophase) or in G1 entry (**Fig. 1e** and **Extended Data** Fig. 1e). Moreover, we did not observe differences in chromosome missegregation between the two conditions in IF staining. These results support that our RAD21 degradation system does not interfere with proper cell cycle progression and enables evaluation of the post-mitotic effects of RAD21 uncoupled from its mitotic role.

### Acute cohesin depletion in G1 recapitulates higher-order 3D chromatin architecture changes seen in asynchronous perturbation

Cohesin depletion in asynchronous cells was shown to be critical for the maintenance of the hierarchical 3D chromatin organization observed in interphase leading to a nearly complete loss of TADs and loops (mostly structural), while reciprocally strengthening compartmentalization.^33,34,38^ Importantly, many of these architectural features are naturally perturbed during mitosis and need to be re-established during the M-to-G1 transition along with the reactivation of the transcriptional program.^5–8,10^ To examine the degree to which cohesin is required for proper post-mitotic reorganization of the 3D architecture, we treated *Rad21*-dTAG cells either with dTAG or DMSO throughout the M-to-G1 transition using our mitotic arrest and release method described above and performed Micro-C analysis (**Fig. 1b**). RAD21 depletion during the M-to-G1 transition prevented the re-formation of TADs globally, which were drastically weaker compared to the DMSO control (**Fig. 1f**). On the other hand, RAD21-depleted G1 cells formed stronger compartments across all chromosomes, supporting the previously reported competition between cohesin-mediated loop extrusion and compartmentalization (**Fig. 1g, h**). In further support, quantification of the contact probabilities across genomic distances upon RAD21 depletion showed reduced interactions at 50 kb–2 Mb ranges and increased interactions beyond 2 Mb, consistent with weakened TADs and strengthened compartmentalization (**Extended Data** Fig. 1f). Importantly, comparison of our results to previously published Micro-C data from asynchronous mESCs with acute RAD21 depletion via auxin-inducible degron (AID)^33^ demonstrated very similar effects of RAD21 depletion on both TADs and compartments between G1 and asynchronous cells, suggesting that both re-establishment and maintenance of these structures are equally dependent on (for TADs) or counteracted by (for compartments) cohesin (**Fig. 1f, g** and **Extended Data** Fig. 1f).^38^

### Regulatory loops with active chromatin features can reform upon mitotic exit in a cohesin- independent manner

To investigate changes in chromatin looping, we called chromatin interactions (loops) at 5 kb, 10 kb, and 25 kb resolution at each sample using Chromosight^51^, the computer vision-based loop calling algorithm, and assembled a list of 64,841 union loops across all samples (see methods) (**Fig. 2a**). Upon RAD21 depletion, over 85% of the union loops were not properly re-established during the G1 phase showing substantially weaker loop strengths (DOWN group, Δ loop score < −0.2) compared to DMSO controls (**Fig. 2b**). The rest of the loops were insensitive to cohesin depletion, with about 14% remaining unaffected (NO group, |Δ loop score| < 0.2), while a minor fraction (0.3%) showed increased strength (UP group, Δ loop score > 0.2) (**Fig. 2b** and **Extended Data** Fig. 2a). Enrichment analysis of accessible regions at the loop anchors, performed using Locus Overlap Analysis (LOLA)^52^, showed significant enrichment for CTCF and cohesin complex (RAD21, SMC1A, and SMC3) at the RAD21-dependent loops (DOWN group) as expected (**Fig. 2c**). Proteins enriched at the RAD21- independent loops (UP/NO group) included components of the Mediator and RNA polymerase complex, ESC-related master regulators, and epigenetic modifiers mostly associated with transcriptional activation. There was also a strong enrichment for YY1, a transcription factor (TF) with reported cohesin-independent architectural function mediating E-P interactions.^33,53–55^ These features strongly suggest that RAD21-independent loops (UP/NO group) likely involve active regulatory elements. To further test this, we categorized all Micro-C loop anchors into four categories: promoters (P; overlap with TSS and H3K4me3 peak), enhancers (E; overlap with H3K27ac peak), structural (S; overlap with RAD21 or CTCF peak), and X (no overlap with any of these features) (**Fig. 2a**). Although >90% of structural loops (S-S, S-X; 31,111 out of 34,134) failed to properly reform during the M-to-G1 transition upon RAD21 depletion (**Fig. 2d, e**), more than 35% of the regulatory loops (P-P, P-E, E-E; 2,021 out of 5,609) were re-established upon mitotic exit at levels similar to the DMSO control, as shown both by scatterplots of loop scores and APA plots (**Fig. 2f, g**). Among the structural loops, we observed that longer loops showed proportionally stronger dependency on RAD21 (up to 1 Mb distance), in agreement with previous studies and the loop-extrusion role of cohesin (**Fig. 2h**).^26–29,32^ This distance effect was stronger in structural loops than in hybrid (regulatory-non regulatory) loops and was not evident in regulatory interactions, which displayed relatively consistent perturbation levels across distances.

**Fig. 2.**
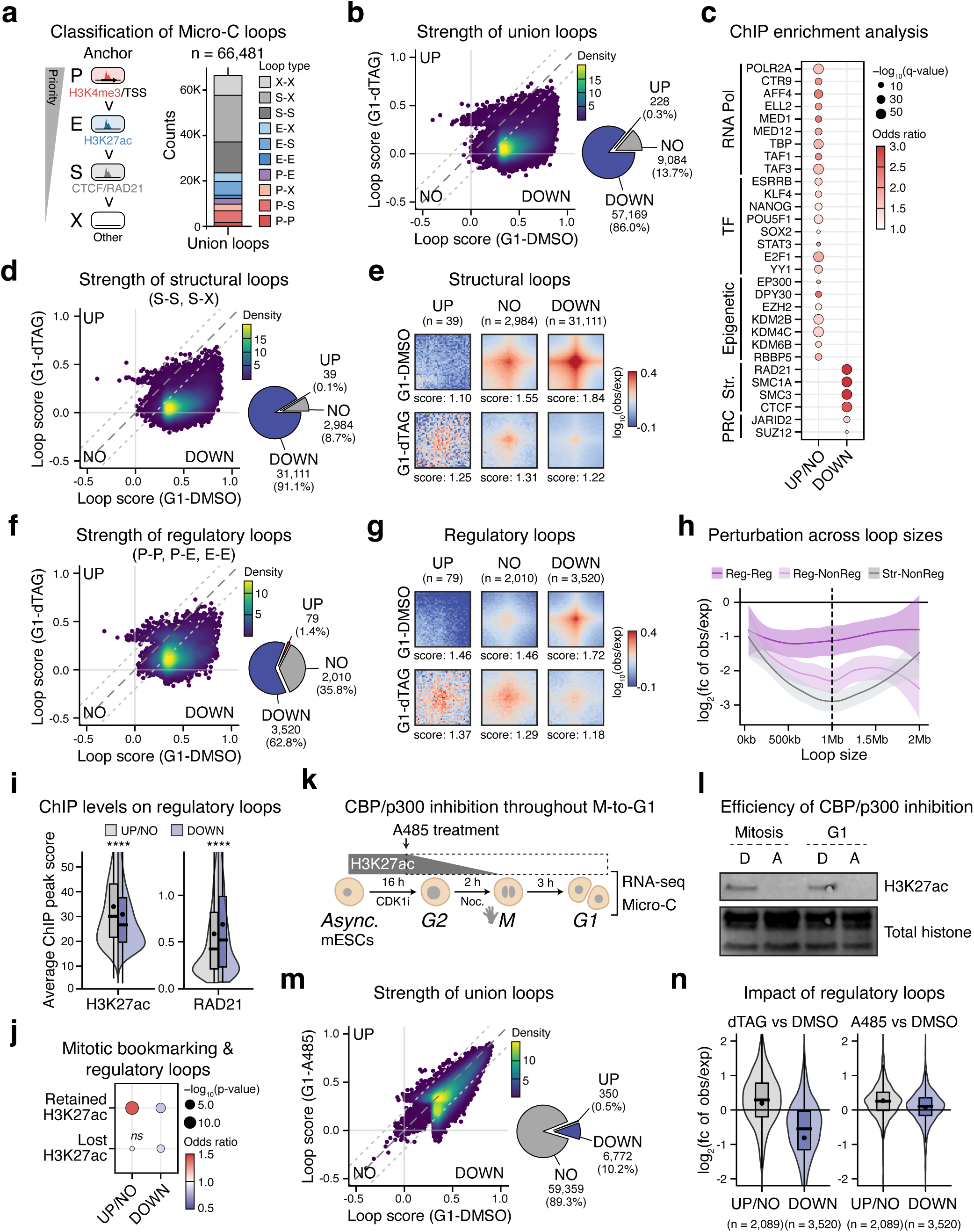
| Post-mitotic resetting of regulatory loops with active chromatin landscape is resistant to acute RAD21 depletion in mESCs. **a**, Micro-C analysis identified a total of 66,481 union loops across all resolutions (5 kb, 10 kb, 25 kb) and datasets. Loop anchors were classified as Promoter (P), Enhancer (E), Structure (S), or X anchors based on hierarchical criteria: P (TSS + H3K4me3), E (H3K27ac), S (RAD21/CTCF), and X (absence of these features). In this study, S-S and S-X loops are categorized as “structural loops”, while P-P, P-E, and E-E loops are categorized as “regulatory loops”. **b**, Left panel: Scatterplot comparing loop strength (Chromosight score) of union loops between G1- DMSO and G1-dTAG Micro-C data. A difference cutoff of 0.2 was applied to classify loops as UP, NO, or DOWN change upon RAD21 depletion. Color represents the density of dots. Right panel: Pie chart illustrating the proportion and number of loops in each category. **c**, Dot plot showing top enriched features for UP/NO or DOWN loops (as defined in **panel b**) based on ChIP enrichment analysis using the LOLA algorithm at the respective loop anchors. The analysis was limited to accessible regions defined by ATAC-seq peaks^61^ within union loop anchors. Only features with significant enrichment in either loop category (q-value < 0.05) are displayed. Dot size represents −log_10_(q-value), while color represents the odds ratio of enrichment. A complete list of significant results is provided in **Supplementary Table 5**. **d**, Left panel: Scatterplot comparing loop strength (Chromosight score) of structural loops (S-S, S-X) between G1-DMSO and G1-dTAG Micro-C data. A difference cutoff of 0.2 was applied to classify loops as UP, NO, or DOWN change upon RAD21 depletion. Color represents the density of dots. Right panel: Pie chart illustrating the proportion and number of loops in each category. **e**, Aggregate peak analysis (APA) of structural loops (S-S, S-X) categorized into UP, NO, or DOWN groups (as defined in **panel d**) in G1-DMSO or G1-dTAG. The number of loops in each group is shown in parentheses. The scores at the bottom represent the ratio of the signal intensity at the center to the signal intensity at the corner. Color represents log_10_ of observed (obs) contact frequency over expected (exp) contact frequency. **f**, Left panel: Scatterplot comparing loop strength (Chromosight score) of regulatory loops (P-P, P-E, E-E between G1-DMSO and G1-dTAG Micro-C data. A difference cutoff of 0.2 was applied to classify loops as UP, NO, or DOWN change upon RAD21 depletion. Right panel: Pie chart illustrating the proportion and number of loops in each category. **g**, APA of regulatory loops (P-P, P-E, E-E) categorized into UP, NO, or DOWN groups (as defined in **panel f**) in G1-DMSO or G1-dTAG. The number of loops in each group is shown in parentheses. The scores at the bottom represent the ratio of the signal intensity at the center to the signal intensity at the corner. Color represents log_10_ of observed (obs) contact frequency over expected (exp) contact frequency. **h**, Line plot showing the average log_2_ fold change of observed (obs) contact frequency over expected (exp) contact frequency across different loop sizes. The ribbon represents the standard error of the mean. “Reg” refers to P or E anchors, “NonReg” refers to S or X anchors, and “Str” refers to S anchors. **i**, Violin and box plot showing the distribution of H3K27ac and RAD21 ChIP-seq/exo peak scores at the anchors of regulatory loops (P-P, P-E, E-E) that are weakened (DOWN) or not (UP/NO) upon RAD21 depletion (as defined in **panel f**). Boxes represent the interquartile range, lines indicate the median, and dots represent the mean. Statistical significance was assessed using the Wilcoxon test (****p < 0.0001). **j**, Dot plot showing the relative enrichment significance of mitotically bookmarked (retained) and non-bookmarked (lost) H3K27ac ChIP-seq peaks between RAD21- dependent (DOWN) and independent (UP/NO) regulatory loops (as defined in **panel f**). Statistical significance was assessed using Fisher’s exact test (*ns* = not significant). Dot size represents −log_10_(p-value), while color represents the odds ratio of enrichment. **k**, Experimental strategy for CBP/p300 inhibition using A485 during the M-to-G1 transition. ESCs were synchronized in mitosis by treatment with 9 µM CDK1 inhibitor for 16 hours, followed by 100 ng/mL nocodazole treatment for 2 hours. After mitotic shake-off, cells were released into G1 for 3 hours. To ensure CBP/p300 inhibition and loss of H3K27ac upon G1 entry, cells were treated with 20 µM A485 starting at the last hour of CDK1 inhibitor treatment, with 0.04% v/v DMSO used as a control. Cells in G1 were collected for RNA-seq and Micro-C. **l**, Representative Western blot showing loss of H3K27ac upon CBP/p300 inhibition in both mitosis and G1 following A485 treatment. DMSO (D). A485 (A). **m**, Left panel: Scatterplot comparing loop strength (Chromosight score) of union loops between G1-DMSO and G1- A485 Micro-C data. A difference cutoff of 0.2 was applied to classify loops as UP, NO, or DOWN change upon A485 depletion. Color represents the density of dots. Right panel: Pie chart illustrating the proportion and number of loops in each category. **n**, Violin and box plot showing the relative changes of RAD21-dependent (DOWN) or independent (UP/NO) regulatory loops (as defined in **panel f**) in the context of RAD21 depletion (dTAG vs DMSO) or upon CBP/p300 inhibition (A485 vs DMSO). The distribution of log_2_ fold change of observed (obs) contact frequency over expected (exp) contact frequency is plotted. Boxes represent the interquartile range, lines indicate the median, and dots represent the mean. The number of loops in each group is shown in parentheses.

Focusing on the regulatory loops, we observed that the RAD21-dependent loops (DOWN group) were strongly enriched for CTCF and core cohesin subunits binding (**Fig. 2i** and **Extended Data** Fig. 2b, c). Intriguingly, the cohesin-loading factor NIPBL was enriched on the RAD21-independent regulatory loops (UP/NO group), suggesting additional cohesin-independent regulatory roles for NIPBL (**Extended Data** Fig. 2c).^56–59^ Overall, RAD21-independent regulatory loop anchors (UP/NO group) were preferentially enriched for occupancy by the transcriptional machinery and CBP/p300, and, accordingly, had significantly stronger ChIP-seq signals for the active promoter and enhancer marks, H3K4me3 and H3K27ac (**Fig. 2i** and **Extended Data** Fig. 2b, c). Furthermore, RAD21- independent loops (DOWN group) showed a significant association with previously defined ESC superenhancers^13,60^ (**Extended Data** Fig. 2d, e) and 3D hyperconnected hubs^13,61^ (**Extended Data** Fig. 2f), suggesting that post-mitotic re-formation of these interactions occurs largely independent of RAD21.

Previous findings from our lab and others have shown that mitotic retention of select transcription factors (e.g. OCT4, SOX2, ESRRB, KLF4, and TBP) and histone marks (H3K27ac), a phenomenon called “mitotic bookmarking”, facilitates rapid and faithful post-mitotic transcription activation of pluripotency-related genes and enhancers in mouse ESCs.^8,40,42,62–65^ As RAD21- independent loops showed a strong enrichment for binding of many of these factors and marks in asynchronous mouse ESCs, we then systematically tested the relative degree of association with their mitotically retained versus lost binding sites. Fisher’s exact test indicated that RAD21- independent loops (UP/NO group) had a preferential enrichment for the H3K27ac bookmarked compared to lost sites (**Fig. 2j**), while no selective enrichment was observed for bookmarking by any other factors (**Extended Data** Fig. 2g).

The abovementioned association analyses suggest that active regulatory features, including H3K27ac bookmarking, might play important roles in the post-mitotic re-assembly of regulatory loops independently of RAD21. However, mitotic inhibition of CBP/p300 with A485, a chemical inhibitor of CBP/p300, which deposits H3K27ac, has been reported to affect only post-mitotic transcriptional activation and not 3D reorganization, previously assessed by Hi-C.^8^ Given that Hi-C is not as sensitive in detecting regulatory interactions, we decided to revisit this question by applying Micro-C. Moreover, we also extended the duration of CBP/p300 inhibition throughout the M-to-G1 transition (mimicking the RAD21 depletion conditions) (**Fig. 2k**). This resulted in near complete loss of H3K27ac both in mitosis and G1 (**Fig. 2l**) with successful mitotic release into G1 both validated with FACS (**Extended Data** Fig. 3a) and H3S10ph staining. Consistent with the previous finding, we observed dramatic perturbation of post-mitotic transcriptional activation (**Extended Data** Fig. 3b), while the 3D reorganization was only mildly perturbed (**Extended Data** Fig. 3c–f). Specifically, we observed a moderate weakening of TADs (**Extended Data** Fig. 3d) and compartments (particularly of A compartments) (**Extended Data** Fig. 3e, f) across the genome. Over 90% of union loops reformed at levels similar to the DMSO control despite the continuous CBP/p300 inhibition (**Fig. 2m**). Importantly, RAD21-independent regulatory loops (UP/NO group), which were enriched in H3K27ac signal and bookmarking, did not show any preferential sensitivity to CBP/p300 inhibition suggesting that CBP/p300 (or H3K27ac) alone cannot explain the efficient post-mitotic re-formation of these interactions (**Fig. 2n**).

Collectively, these findings highlight that regulatory loops associated with strong enhancers and transcriptional activity can be effectively re-assembled upon mitotic exit in the absence of RAD21, suggesting they are driven by cohesin-independent mechanisms. Importantly, inhibition of CBP/p300 activity and consequently impaired transcriptional activity had also minimal effects on the re-formation of these regulatory interactions, arguing for the presence of complex and likely redundant compensatory mechanisms—such as the abundance of other active histone marks and protein factors that could drive clustering and condensation.

### Post-mitotic gene reactivation is largely independent of cohesin with few exceptions in mESCs

Previous studies in various asynchronous cell lines have reported a rather limited impact of acute cohesin-degradation on the global transcriptional activity in contrast to the widespread and dramatic effects on 3D chromatin architecture. A potential explanation for this long-standing puzzle was that ongoing transcriptional activity can be sustained in the short term by cohesin-independent regulatory interactions. However, it is plausible that transcriptional reactivation upon mitotic exit is more vulnerable to cohesin degradation than steady-state interphase transcriptional activity, when all regulatory factors, transcriptional machinery, and loops are re-assembled. To test this hypothesis, we collected mouse ESCs (*Rad21*-dTAG cells) at G1 after treatment with dTAG or DMSO throughout the M-to-G1 transition, as described before (**Fig. 1b**). RNA-seq analysis revealed that only a small subset of genes showed impaired reactivation upon RAD21 degradation (**Fig. 3a**). Importantly, the extent of transcriptional perturbations in G1 (**Fig. 3a**) was similar to the ones reported in asynchronous mouse ESCs^33,66^, arguing that both maintenance and post-mitotic reactivation of transcription in ESCs are largely independent of cohesin with only a few dozen genes being affected.

**Fig. 3.**
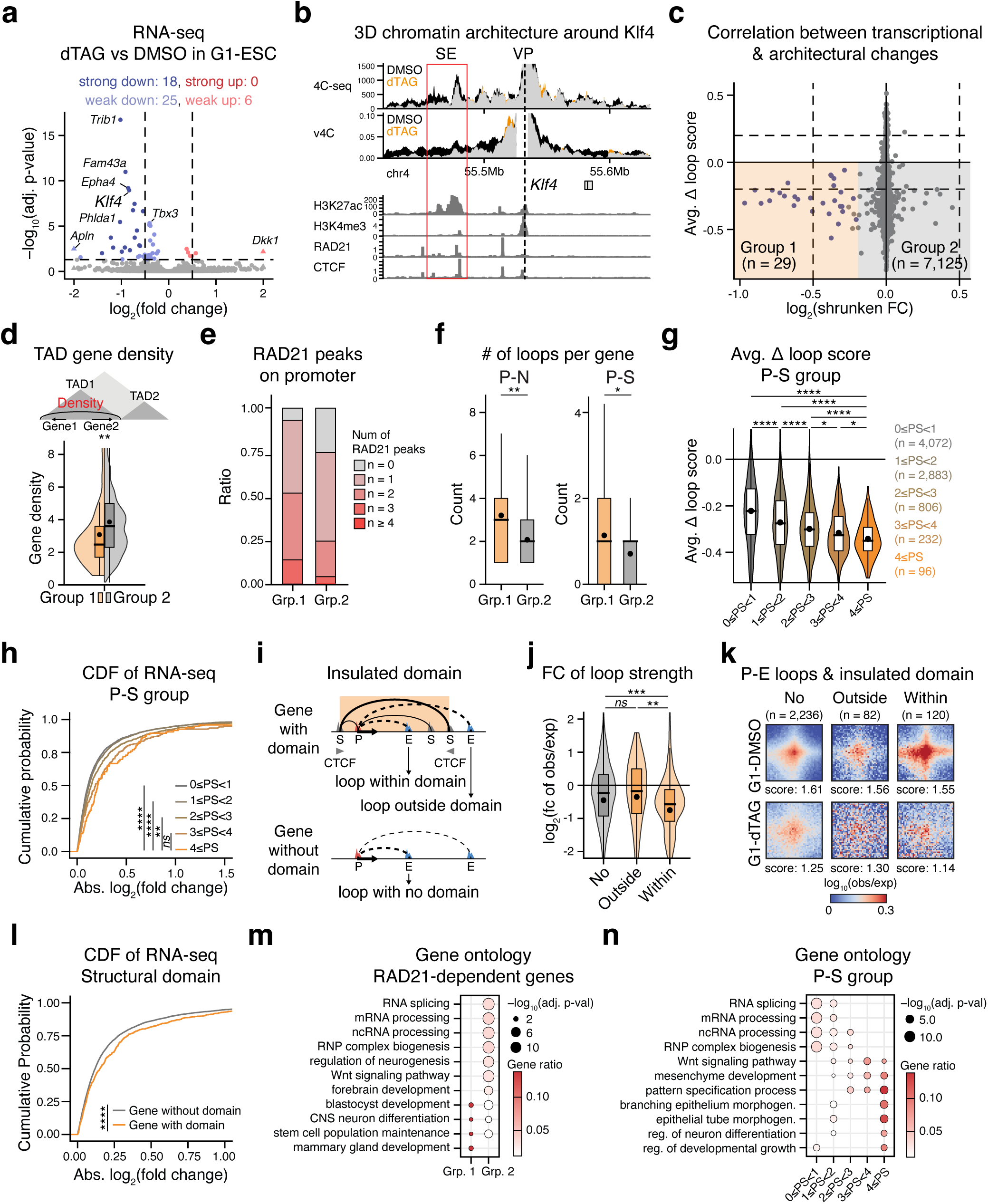
| Post-mitotic gene reactivation is largely independent of cohesin with few exceptions in mESCs. **a**, Volcano plot of RNA-seq data highlighting transcriptional changes upon dTAG treatment in G1-ESC. Statistical significance was determined using an adjusted p-value cutoff of 0,05, and changes were classified as strong or weak based on an absolute shrunken log_2_ fold change cutoff of 0.5. The analysis was performed using n=3 replicates for DMSO and n=5 replicates for dTAG. **b**, Top panel: Experimental 4C-seq data analysis of ESCs in G1 treated with DMSO or dTAG throughout the M-to-G1 transition, shown as average CPM around the viewpoint (VP, *Klf4* promoter). Regions with higher 4C-seq signals in DMSO are shown in black, while those with higher signals in dTAG are shown in orange. Middle panel: Virtual 4C calculated from Micro-C of ESCs in G1 with DMSO or dTAG treatment, represented similarly. Bottom panel: ChIP-seq (H3K27ac) and ChIP-exo (H3K4me3, RAD21, and CTCF) tracks. The superenhancer (SE) upstream of the *Klf4* promoter is highlighted in red. **c**, Scatterplot showing the correlation between transcriptional and architectural changes upon RAD21 depletion during the M-to-G1 transition. The x-axis shows the log_2_ shrunken fold change (FC) in RNA-seq (dTAG vs DMSO in G1-ESC), while the y-axis shows the average difference (Δ) in loop scores for all promoter-originating loops (P-P, P-E, P-S, P-X) per gene. Only genes detected in RNA- seq and have promoter-originating loops are plotted. Group 1 (n = 29, highlighted in orange) includes all significantly downregulated genes upon dTAG treatment that show a concordant decrease in average loop strength. Group 2 (n = 7,125, highlighted in grey) includes all genes that also show a decrease in average loop strength but without significant transcriptional changes. **d**, Box plot and violin plot showing the distribution of gene density within TADs where genes in Group 1 (orange) or Group 2 (grey) genes are located. Boxes represent the interquartile range, lines indicate the median, and dots represent the mean. Statistical significance was assessed using the Wilcoxon test (**p < 0.01). **e**, Stacked bar plot showing the proportions of Group1 or Group2 genes with different number of RAD21 ChIP-exo peaks on their promoters (−/+ 5 kb from TSS). **f**, Box plots showing the significant differences either in the overall promoter connectivity (number of P-N loops, N: any type of anchor) of Group 1 and Group 2 genes (left panel) or specifically of the numbers of P-S (right panel). Boxes represent the interquartile range, lines indicate the median, and dots represent the mean. Statistical significance was assessed using the Wilcoxon test (*p < 0.05, **p < 0.01). **g**, Box plot and violin plot showing the average difference (Δ) in loop scores for all promoter-originating loops per gene, categorized based on the number of P-S loops they form. The number of loops in each category is shown in parentheses. Boxes represent the interquartile range, lines indicate the median, and dots represent the mean. Statistical significance was assessed using the Wilcoxon test (*p < 0.05, ****p < 0.0001). **h**, Cumulative distribution function (CDF) plot showing the absolute log_2_ fold change of RNA-seq (dTAG vs DMSO in G1-ESC), measuring the overall transcriptional perturbation upon RAD21 depletion during the M-to-G1 transition. Genes were grouped based on the number of P-S loops (as defined in **panel g**). Statistical significance was assessed using a two-tailed, two-sample Kolmogorov-Smirnov test (*ns* = not significant, **p < 0.01, ****p < 0.0001). **i**, Schematic showing example genes either within an insulated domain (upper panel) or without an insulated domain (bottom panel). An insulated domain is defined as the largest S-S loop encompassing a gene, with convergent CTCF motifs at its anchors. P-E loops are depicted based on whether they fall inside or outside the insulated domain. **j**. Box plot and violin plot showing the log_2_ fold change of observed (obs) contact frequency over expected (exp) contact frequency of loops categorized as No Domain (No), Outside Domain (Outside), and Within Domain (Within), as defined in **panel i**. Boxes represent the interquartile range, lines indicate the median, and dots represent the mean. Statistical significance was assessed using the Wilcoxon test (*ns* = not significant, **p < 0.01, ***p < 0.001). **k**, Aggregate peak analysis (APA) of P-E loops categorized based on insulated domain (as defined in **panel i**) in G1-DMSO or G1-dTAG. The number of loops in each group is shown in parentheses. The scores at the bottom represent the ratio of the signal intensity at the center to the signal intensity at the corner. Color represents log_10_ of observed (obs) contact frequency over expected (exp) contact frequency. **l**, Cumulative distribution function (CDF) plot showing the absolute log_2_ fold change of RNA-seq (dTAG vs DMSO in G1-ESC), measuring the overall transcriptional perturbation upon RAD21 depletion during the M-to-G1 transition. Genes were grouped based on the presence of an insulated domain, as defined in **panel i**. Statistical significance was assessed using a two-tailed, two-sample Kolmogorov-Smirnov test (****p < 0.0001). **m**, Dot plot showing select biological processes significantly enriched in Group 1 or Group 2 genes. Dot size represents −log_10_(adjusted p-value), while color represents the gene ratio of enrichment. An adjusted p-value cutoff of 0.05 and a q-value cutoff of 0.2 were used to determine significance. A complete list of significant results is provided in **Supplementary Table 6**. **n**, Dot plot showing select biological processes significantly enriched in gene categorized based on the number of P-S loops they from. Dot size represents −log_10_(adjusted p- value), while color represents the gene ratio of enrichment. An adjusted p-value cutoff of 0.05 and a q-value cutoff of 0.2 were used to determine significance. A complete list of significant results is provided in **Supplementary Table 6**.

To understand the preferential vulnerability of some genes on RAD21 depletion, we focused on *Klf4*, an important ESC-related gene with high transcriptional sensitivity to RAD21 degradation both in asynchronous cells^33,48^ and during the M-to-G1 transition (our study) (**Fig. 3a**). Importantly, using a KLF4-MS2 transcriptional reporter in the context of the *Rad21*-dTAG cell line (**Extended Data** Fig. 4a), we confirmed through real-time, single-cell imaging that the efficiency of transcriptional reactivation of *Klf4* in cells exiting mitosis was significantly lower in the absence of RAD21 (**Extended Data** Fig. 4b–e). Previous high-resolution single-cell imaging showed that *Klf4* nascent transcription activity shows bursts that are controlled by interactions with a distal cluster of enhancers, also identified as superenhancer^60^, located about 50–65 kb downstream.^48^ The observation that RAD21 depletion resulted in a smaller fraction of *Klf4* alleles showing bursts during the M-to-G1 reactivation indicates that RAD21 depletion affects the probability of a *Klf4* allele turning on and/or its bursting frequency once turned on (**Extended Data** Fig. 4e). At the same time, burst amplitude and duration were not affected by RAD21 depletion (**Extended Data** Fig. 4f). These results resemble the reduction in E-P proximity and number of busting alleles/burst frequency upon RAD21 depletion previously seen in asynchronous cells and suggest impaired *Klf4* P-E interaction during the reactivation process.^48^ Consistent with this notion, our Micro-C analysis on G1 cells detected a substantially weaker interaction between the *Klf4* promoter and the 50-65 kb downstream superenhancer upon RAD21 depletion (**Fig. 3b**). This was also true for *Tbx3*, another ESC-associated gene sensitive to RAD21 depletion (**Extended Data** Fig. 5a). High-resolution 4C-seq experiments showed high correlation with the Micro-C data and independently validated the perturbed P-E interactions around *Klf4* and *Tbx3* (**Fig. 3b** and **Extended Data** Fig. 5a). These results suggest a mechanistic link between disrupted regulatory contacts and transcriptional perturbation upon cohesin-degradation during the M-to-G1 reactivation, in agreement with previous results from high-resolution single-gene imaging in asynchronous cells.^48^

To determine the molecular basis for the transcriptional vulnerabilities or insensitivity of genes on cohesin degradation, we then tested on a genome-wide scale the concordance between the extent of transcriptional perturbation and the mean architectural reorganization around each gene (average delta score of all promoter-originating loops; P-P, P-E, P-S, P-X) (**Fig. 3c**). Of note, only expressed genes with at least one Micro-C detected promoter-originating loop were included in this analysis. The scatterplot revealed that genes with significantly decreased transcription (Group 1, n = 29, adjusted p-value < 0.05) experienced, on average, weakening of promoter-originating loops, suggesting a correlation between changes in chromatin architecture and transcriptional disruption. However, thousands of other genes within the RAD21-insensitive group (Group2, genes that experienced architectural but not transcriptional changes, n = 7,125) also exhibited disruption of promoter-anchored loops, often to a higher extent. To investigate why Group1 genes exhibit greater transcriptional sensitivity to cohesin loss compared to Group2 genes, we then examined various characteristics, including the nature and number of loops around each gene and several features of their topological neighborhood. Overall, the TAD sizes were comparable between Group 1 and Group 2 genes and the distance to the closest boundaries was also very similar (median of 200 kb) (**Extended Data** Fig. 5b, c). However, Group 1 genes (RAD21-sensitive) were preferentially located near strong boundaries, which exhibited significantly greater loss of insulation upon RAD21 depletion (**Extended Data** Fig. 5d). Moreover, TADs with Group 1 genes (RAD21-sensitive) exhibited significantly lower gene density (**Fig. 3d**), a trend further supported by the H3K4me3 ChIP-exo peak density plot (**Extended Data** Fig. 5e). To assess whether this pattern is generalizable, we ranked TADs based on their density of transcriptionally active genes and divided them into quintiles (**Extended Data** Fig. 5f). Analyzing the cumulative probability of absolute log_2_ fold change from RNA-seq revealed that genes in gene-poor TADs were more transcriptionally perturbed by RAD21 degradation (**Extended Data** Fig. 5g). This suggests that a higher density of nearby active promoters may contribute to greater resilience to RAD21 depletion when it comes to transcriptional reactivation.

Another observation that could explain the preferential sensitivity of Group 1 genes to RAD21 deletion was the prevalence of RAD21 binding on their promoters. Specifically, 93% of genes in Group 1 had at least one RAD21 promoter peak with more than 50% of genes having two or more peaks, while only 76% of Group 2 genes (RAD21-insensitive), were occupied by RAD21 around their TSS (**Fig. 3e**). Also, we observed that Group 1 genes had overall higher promoter connectivity (number of distinct interactions, P-N) compared to Group 2 genes and a significantly higher number of P-S interactions (**Fig. 3f** and **Extended Data** Fig. 5h). To orthogonally validate these findings, we grouped genes based on their P-S loop connectivity. Genes with more P-S loops experienced greater loop-level perturbation upon RAD21 depletion (**Fig. 3g**), and—more importantly—increased transcriptional perturbation (**Fig. 3h**). Together, these data suggest that genes engaged in more RAD21-homotypic loops (RAD21 both on the promoter and the interacting anchor) are more dependent on cohesin.

Cohesin-mediated extrusion, along with the resulting loops and insulated domains, has been shown to promote proximity and interactions between enhancers and promoters actively. Therefore, we hypothesized that genes encompassed within insulated domains^67,68^ might show increased sensitivity upon cohesin depletion. To test this, we focused on genes that are surrounded by S-S loops with convergent CTCF binding sites on their anchors (**Fig. 3i**). For genes encompassed by multiple insulated domains, the largest domain was selected for analysis (genes with domain), while the genes lacking surrounding structural loops (genes without domain) were used as a control. Interestingly, P-E regulatory loops confined within insulated domains exhibited greater perturbation upon RAD21 depletion compared to those located outside the domains or unrelated to any domain (**Fig. 3j, k**). This suggests that the structural integrity of insulated domains actively supports the formation and stability of regulatory loops within them. Furthermore, genes associated with insulated domains showed higher transcriptional disruption upon RAD21 depletion compared to genes without such domains, as demonstrated by the cumulative probability plot (**Fig. 3l**).

Finally, to gain insights into the biological relevance of the RAD21-sensitive genes, we next performed Gene Ontology (GO) analysis. Despite the small number, Group 1 genes showed significant and preferential enrichment for GO categories related to both stem cell maintenance and differentiation (**Fig. 3m**). Moreover, genes with higher P-S connectivity (but not higher P-E or P-P) also displayed a clear enrichment for differentiation-related terms (**Fig. 3n**), suggesting that genes critical for developmental processes might rely on multiple structural loops and cohesin for proper regulation.

Collectively, our findings demonstrate that transcriptional sensitivity to RAD21 depletion is limited and intricately tied to the architectural environment, characterized by promoter-originating structural (P-S) loops and insulated domains. Genes lacking these structural features exhibited transcriptional resilience, which can be attributed to their association with high gene density, and enrichment of housekeeping gene functions, consistent with the well-established P-P network in housekeeping genes.^69^ In contrast, sensitive genes were associated with both stem cell maintenance and differentiation pathways, raising intriguing questions about potential context-specific roles of cohesin during cellular differentiation.

### The transition from naïve to formative pluripotent state is partially dependent on cohesin

The previous results suggested that cohesin plays a limited role in the post-mitotic reactivation of the ESC transcriptional program, and preferentially affects genes involved in developmental processes. Therefore, we hypothesized that cohesin depletion will have more dramatic effects on the activation and establishment of a new developmental program. To test this hypothesis, we synchronized mouse ESCs (*Rad21*-dTAG cells) in mitosis and then released them into G1 in the presence of strong differentiation signals (Activin A, FGF2) in addition to dTAG treatment (or DMSO as control) (**Fig. 4a** and **Extended Data** Fig. 6a). These signals are known to initiate the transition from a naïve pluripotency state (ESCs) to the formative pluripotency state (epiblast-like cells (EpiLCs)).^70,71^ Differential RNA-seq analysis (focusing only on DMSO samples) revealed that the short exposure of cells exiting mitosis to the EpiLC signals was sufficient to induce significant transcriptional changes compared to the ESC (self-renewal) conditions (**Extended Data** Fig. 6b, c). The detected up- and downregulated genes followed the expected trends and signatures reported in previous ESC-to- EpiLC differentiation studies^72–74^ (**Extended Data** Fig. 6d), demonstrating an early *de novo* activation of many EpiLC-related genes and gradual silencing of ESC-related programs. These results validate that acute exposure to differentiation signals during the M-to-G1 transition induces an early transcriptional reprogramming towards an EpiLC state.

**Fig. 4.**
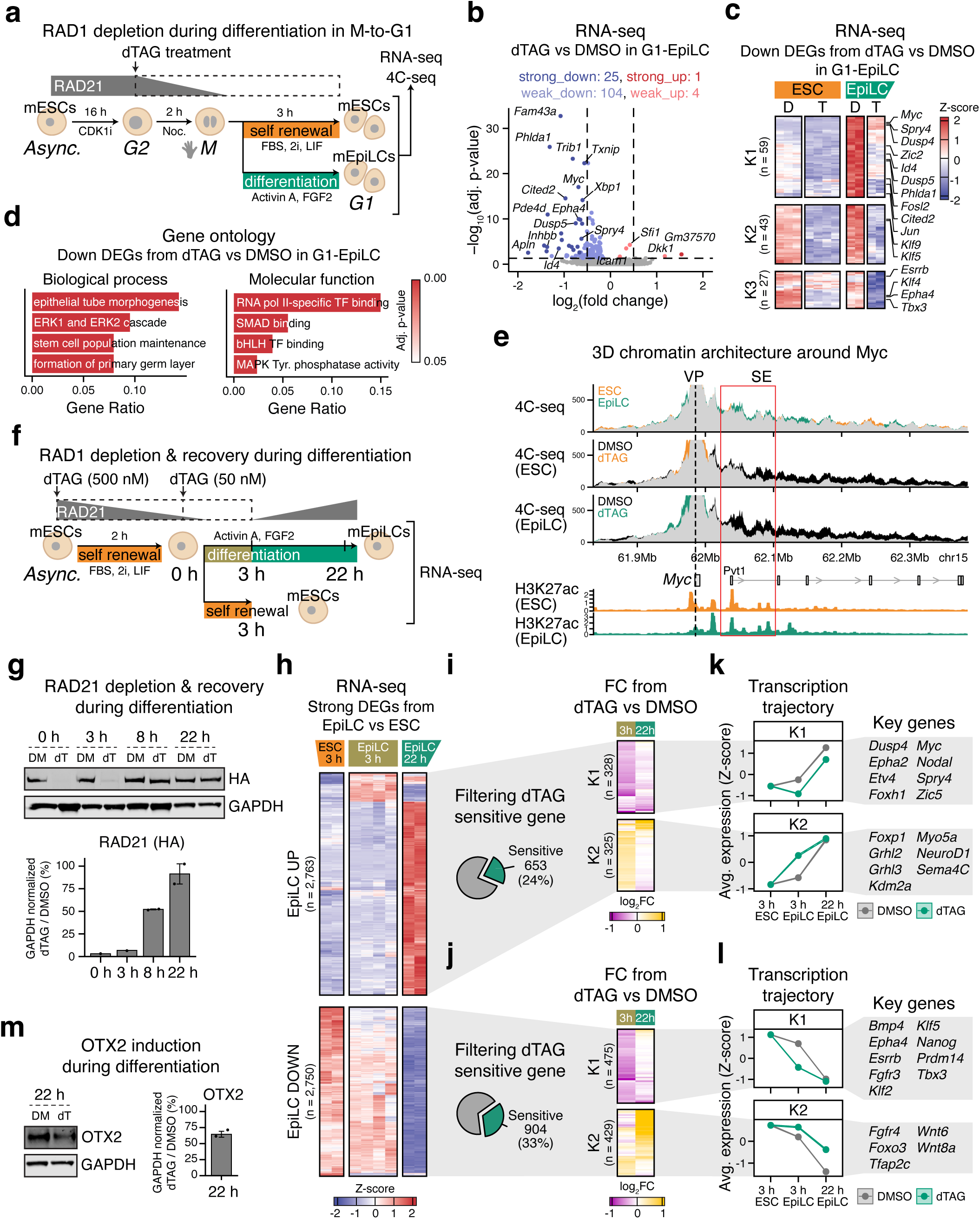
| The transition from naïve to formative pluripotent state is partially dependent on RAD21. a, Experimental strategy for assessing the impact of RAD21 depletion during the M-to-G1 transition under self-renewal (naïve pluripotent) conditions or during acute differentiation into formative pluripotency (epiblast-like cells, EpiLC). After mitotic shake-off, cells are released into G1 for 3 hours under either self-renewal conditions (FBS, 2i, LIF) or differentiation conditions (Activin A, FGF2). To ensure RAD21 depletion upon G1 entry, cells were treated with 500 nM dTAG starting at G2, with 0.05% v/v DMSO used as a control. Cells in G1 were collected for RNA-seq and 4C-seq. **b**, Volcano plot of RNA-seq data highlighting transcriptional changes upon dTAG treatment in G1-EpiLC. Statistical significance was determined using an adjusted p-value cutoff of 0,05, and changes were classified as strong or weak based on an absolute shrunken log_2_ fold change cutoff of 0.5. The analysis was performed using n=2 replicates per each condition. **c**, Heatmap showing the Z-score of RNA-seq values in G1-ESC and G1-EpiLC with DMSO (D) or dTAG (T) treatment. DEGs that are significantly downregulated in dTAG vs DMSO in G1-EpiLC (as shown in **panel b**) are plotted. Genes are clustered into 3 groups with K-means clustering. A list of genes in each group is provided in **Supplementary Table 7**. **d**, Bar plot showing select biological process (left panel) and molecular function (right panel) from enriched in DEGs that are significantly downregulated in dTAG vs DMSO in G1-EpiLC (as shown in **panel b**). Color represents adjusted p-value. An adjusted p-value cutoff of 0.05 and a q-value cutoff of 0.2 were used to determine significance. A complete list of significant results is provided in **Supplementary Table 6**. **e**, Experimental 4C-seq data analysis comparing interaction patterns around the *Myc* promoter viewpoint (VP) between ESC vs EpiLC in DMSO conditions (top panel) or DMSO vs dTAG in ESC conditions (middle panel) or EpiLC conditions (bottom panel). 4C-seq signal is expressed as normalized counts per million (CPM). Regions with higher 4C-seq signals in DMSO are shown in black, while those with higher signals in dTAG are shown in orange. The H3K27ac ChIP- seq tracks for ESC and EpiLC are displayed at the bottom panel. The red box highlights a superenhancer (SE) downstream of the *Myc* promoter, which exhibits increased contacts in EpiLC compared to ESC conditions and undergoes EpiLC-specific perturbation upon dTAG treatment. **f**, Experimental strategy for assessing the impact of transient RAD21 perturbation on ESC-to-EpiLC differentiation. Asynchronous ESCs were treated for 2 hr with 500 nM dTAG under self-renewal conditions, followed by differentiation for 22 hours. During the initial 3 hours of differentiation, 50 nM dTAG was maintained before being washed out for the remainder of the differentiation process. Cells were collected at 0 hour, 3 hours, 8 hours, and 22 hours for Western blot and 3 hours and 22 hours for RNA-seq. Asynchronous cells kept in self-renewal conditions, with or without dTAG treatment, were collected at 3 hours as control. **g**, Top panel: Western blot showing depletion and recovery of RAD21 (HA-tagged) during EpiLC differentiation. Botton: Bar plot showing the quantified intensity of RAD21 relative to GAPDH intensity. The error bar represents the standard error from two biological replicates (**Extended Data** Fig. 7a). **h**, Heatmap showing the trends of strong DEGs (absolute log_2_(fold change) > 0.5) from the EpiLC vs ESC comparisons, shown as Z-scores of RNA-seq across all DMSO conditions (ESC-3h, EpiLC-3h, EpiLC-22h). Genes are clustered using K-means clustering into EpiLC UP (n = 2,763) and EpiLC DOWN (n = 2,750) groups. A list of genes in each group is provided in **Supplementary Table 7**. **i**, **j**, Genes sensitive to dTAG treatment within the EpiLC UP group (n = 635) (**i**) and EpiLC DOWN group (n = 905) (**j**) were selected for downstream analysis. K-means clustering, based on the log₂ fold change of RNA-seq (dTAG vs. DMSO) at 3 hours or 22 hours in EpiLC conditions, identified two subgroups within each category. Color represents log_2_ of fold change from dTAG vs DMSO. A list of genes in each group is provided in **Supplementary Table 7**. **k, l**, Line plot showing the transcriptional trajectory of filtered genes in each cluster, with or without dTAG treatment. The average z-score and its standard error (as ribbon) at each timepoint are plotted. Key genes within each cluster are highlighted on the right panel. **m**, Left panel: Western blot showing a reduced expression of OTX2, a representative marker of the EpiLC state, upon transient RAD21 depletion. Right panel: Bar plot showing the quantified intensity of OTX2 relative to GAPDH intensity. The error bar represents the standard error from two biological replicates (**Extended Data** Fig. 7a)

To determine the role of RAD21 in the *de novo* establishment of the EpiLC program, we then compared the transcriptional outcomes of RAD21 depletion during the M-to-G1 transition in the context of EpiLC compared to ESC condition. Although the extent of transcriptional perturbation was again rather limited, we detected 134 perturbed genes (129 downregulated and 5 upregulated) in G1- EpiLC cells (**Fig. 4b**), compared to only 49 (43 downregulated and 6 upregulated) in G1-ESC cells (**Fig. 3a**). K-means clustering of the downregulated differentially expressed genes (DEGs) upon RAD21 depletion in G1-EpiLC revealed three main categories of perturbation (**Fig. 4c**). The largest group (K1, n = 59) consists of genes that are *de novo* activated during EpiLC differentiation but showed an impaired upregulation upon RAD21 depletion (e.g., *Myc*, *Jun*, *Spry4*, *Dusp4*, *Zic2*, *Phlda1*, and *Fosl2*). The K2 cluster (n = 43) includes genes that are also upregulated in EpiLC conditions compared to their basal level expression in ESC conditions and failed to re-activate in both conditions upon RAD21 depletion (e.g. *Klf5*, *Klf9*). Finally, the K3 cluster (n = 27) contains stem-cell-associated genes such as *Klf4*, *Tbx3*, *Esrrb*, and *Epha4* which are downregulated during EpiLC differentiation and are transcriptionally susceptible to RAD21 depletion. Therefore, ∼80% of genes (K1 and K2 clusters) with impaired transcriptional activity upon cohesin depletion are genes that need to be upregulated during EpiLC differentiation. As expected, GO analysis of all downregulated DEGs upon RAD21 depletion in G1-EpiLC showed a significant association with stem cell maintenance (from K3 cluster) and developmental processes (from K1 & K2 cluster) (**Fig. 4d**). Moreover, downregulated DEGs were strongly enriched for transcription factor genes (40 out of 129 down DEGs were genes coding either TF or TF-binding proteins), suggesting that the activation of TF genes critical for initiating differentiation shows a preferential dependency on cohesin (**Fig. 4d**).

To investigate how RAD21 depletion impairs the activation of EpiLC-related genes, we performed 4C-seq on representative TF genes, *Myc* (**Fig. 4e**) and *Fosl2* (**Extended Data** Fig. 6e), from the K1 cluster under four conditions: ESCs or EpiLCs in G1, treated with either DMSO or dTAG. Comparison of the 4C-seq profiles in ESC and EpiLC conditions revealed that both *Myc* and *Fosl2* gene promoters establish new or stronger interactions with putative EpiLC-specific enhancers—located ∼50 kb downstream for *Myc* and ∼450 kb upstream for *Fosl2*—upon acute exposure to EpiLC signals. RAD21 depletion induced an extensive abrogation of distal interactions in both ESC and EpiLC conditions but also a preferential weakening of the EpiLC-specific P-E interactions. These results suggest that RAD21 is critical for establishing regulatory interactions around developmental genes to support their transcriptional activation during cell fate transitions.

Knowing that early activation of EpiLC-related genes, including critical TFs, exhibits a unique vulnerability to RAD21 depletion, we next sought to determine how these transcriptional perturbations affect ESC-to-EpiLC differentiation. Since cells cannot tolerate prolonged RAD21 depletion over one full cell cycle (∼12 hours in mouse ESCs), we treated asynchronous ESCs with dTAG (or DMSO as control) for 2 hours to degrade RAD21 prior to exposure to differentiation signals (Activin A, FGF2) for an additional 22 hours, after which cells were collected for RNA-seq (**Fig. 4f**). To ensure continued RAD21 depletion during the early stages of differentiation, we maintained cells in low dTAG concentration for the first 3 hours of differentiation, after which the dTAG was washed out to allow RAD21 recovery. Western blot confirmed the nearly complete degradation (∼95%) of RAD21 for up to 3 hours after the initiation of differentiation, followed by a gradual recovery after dTAG washout, reaching ∼60% of normal levels by 8 hours and ∼90% by 22 hours (**Fig. 4g** and **Extended Data** Fig. 7a). The depletion and recovery of RAD21 was also reflected in the cell cycle profiles measured with FACS using DAPI staining (**Extended Data** Fig. 7b). At the 0-hour time point (which is after 2 hours of RAD21 depletion) there was minimal impact on the cell cycle, consistent with our earlier findings. With prolonged dTAG treatment, cells accumulated in the G2/M phase; however, the restoration of RAD21 levels enabled mitotic progression, resulting in a nearly normal cell cycle profile.

We then focused on the transcriptional changes induced by transient RAD21 depletion. Principal component analysis (PCA) separated the samples primarily based on the timepoint of the EpiLC differentiation (3 hours vs 22 hours), while DMSO and dTAG samples from each timepoint were clustered closely together (**Extended Data** Fig. 7c). Differential expression analysis of the DMSO samples revealed >800 strong DEGs induced by the differentiation conditions at the 3-hour timepoint and >5,000 by 22 hours (**Extended Data** Fig. 7d, e), documenting an extensive transcriptional rewiring during ESC-to-EpiLC differentiation, in agreement to previous studies.^72–75^ To understand to what degree transient RAD21 depletion affects the EpiLC transcriptional response, we focused on EpiLC-relevant or ESC-relevant genes, meaning genes that are upregulated (EpiLC UP, n = 2,763) or downregulated (EpiLC DOWN, n = 2,750), respectively, during this transition (**Fig. 4h**). While the majority of these genes (76% of EpiLC UP and 67% of EpiLC down) were irresponsive to RAD21 depletion, K-means clustering of the remaining ones (653 out of 2,763 for EpiLC UP (**Fig. 4i**), 904 out of 2,750 for EpiLC DOWN (**Fig. 4j**)) revealed distinct patterns of dTAG-induced dysregulation, and a strong association with genes and signaling pathways linked to differentiation and development (**Extended Data** Fig. 7f–i). Both for EpiLC UP and DOWN genes, we identified clusters (UP-K2 and DOWN-K1) where transcriptional dysregulation was significant at the 3-hour timepoint but almost fully alleviated by 22 hours when RAD21 level was restored (**Fig. 4k, l**). However, we also detected gene clusters (UP-K1 and DOWN-K2) that initially failed to be upregulated or downregulated at the 3- hour time point and remained persistently dysregulated at 22 hours, despite the recovery of RAD21 levels (**Fig. 4k, l**). The DOWN-K2 cluster involved key developmental regulators and signaling pathway components, such as *Fgfr4*, *Foxo3*, and *Wnt6*, which failed to be properly downregulated by 22 hours following transient dTAG treatment. On the other hand, the UP-K1 cluster comprised important EpiLC-associated genes, including *Epha2*, *Myc*, and *Zic5*, which exhibited impaired upregulation at 22 hours. These effects—although moderate— suggest that transient RAD21 degradation could impair ESC-to-EpiLC differentiation. In agreement, Western blot analysis showed reduced protein levels of OTX2, a *bona fide* marker of the EpiLC state, at 22 hours following differentiation with acute RAD21 depletion (**Fig. 4m**).

In conclusion, our findings in G1 and asynchronous cells demonstrate that acute RAD21 depletion at the onset of ESC-to-EpiLC differentiation induces limited but important transcriptional perturbations which are more extensive than the ones induced in the self-renewal conditions and can have long-lasting effects on cell fate transition independent of cell cycle defects.

## Discussion

Cohesin plays a critical role in shaping 3D chromatin architecture through loop extrusion, facilitating the interaction of distal genomic elements, such as enhancers and promoters, by bringing them into close proximity.^26–29,67,68^ However, acute cohesin perturbation in asynchronous cells was shown to have a limited impact on transcriptional activity.^31–39^ To explain these puzzling findings, one plausible model proposed that both enhancer-promoter (E-P) interactions and transcriptional activity can persist for short periods in the absence of cohesin through “buffering” mechanisms mediated by the presence of chromatin marks and factors.^33^ However, prolonged depletion of cohesin or additional perturbations such as cell division or cell fate transition are expected to have more drastic effects.^33^ Here, we directly tested this model by designing a strategy that enabled acute cohesin depletion in naïve mouse ESCs during cell division (specifically during the M-to-G1 transition) both under self-renewing conditions (maintenance of cell identity) or upon acute differentiation towards a formative EpiLC state (cell fate transition). This approach allowed us to dissect the role of cohesin both on the post-mitotic re-establishment of cell-type specific loops and transcription and also on the *de novo* establishment of loops and programs associated with a new fate. Our results support an overall moderate role of cohesin-mediated extrusion in either setting but also expose locus-specific and context-specific vulnerabilities to cohesin loss and potential compensatory mechanisms.

Unlike the well-established role of cohesin in maintaining 3D chromatin organization, its role in the re-establishment of loops, TADs, and compartments, after their collapse during mitosis due to the drastic condensation and unique folding of the genome, has yet to be resolved.^5^ The essential role of cohesin in cell division makes it hard to perturb it during mitotic exit without affecting the cell cycle progression. To overcome this challenge, instead of perturbing the cohesin complex directly, recent preprint studies in different cell types depleted NIPBL, an accessory factor of the cohesin complex, associated with the loading and processivity of extrusive cohesin.^76–78^ In our study, we were able to directly deplete the core cohesin subunit RAD21 by timing its degradation, ensuring that around 10% of the remaining protein at prometaphase was sufficient to support successful mitotic exit before complete depletion during the M-to-G1 transition. Our Micro-C analysis revealed that acute RAD21 depletion during the M-to-G1 transition drastically impairs the post-mitotic re- formation of TADs and the vast majority (>85%) of the chromatin loops, while strengthening G1 compartments, in agreement with the proposed competing forces of extrusion and compartmentalization.^34,38,79^ In agreement with previous studies, we found that cohesin/CTCF-bound structural interactions are more vulnerable when bridging more distal genomic regions,^39,80,81^ highlighting the requirement of loop extrusion for the formation of long-range interactions. However, regulatory interactions involving promoter or enhancer elements were less affected by cohesion depletion, with 35% being able to reform after mitosis at expected or higher frequencies. The most resilient 3D contacts involved regulatory elements typical of strong, active enhancers and promoters, bound by many TFs and cofactors and decorated by histone acetylation marks, in agreement with findings in asynchronous cells^33,39^ and further supporting the presence of cohesin-independent mechanisms of E-P interactions.^20,39^ Intriguingly, many of the factors enriched in the resilient loops are dissociated from the condensed mitotic chromatin but select marks, such as H3K27ac, are retained and serve as bookmarks critical for rapid reactivation of cell-type specific transcriptional programs.^8^ It is plausible that these bookmarks, upon mitotic exit, facilitate rapid recruitment and re- assembly of transcriptional components, which through multivalent interactions then enable self-organization/micro-compartmentalization and spatial proximity among enhancers and promoters in the absence of cohesin. To test whether the cohesin-independent loops are indeed dependent on bookmarking and H3K27ac, we inhibited CBP/p300 activity throughout the M-to-G1 transition. In addition to wiping off H3K27ac, CBP/p300 inhibition likely perturbs globally the active chromatin state, as this enzyme acetylates and activates many transcription factors and cofactors.^82^ In agreement with its transcriptional role, we observed an extensive impairment of post-mitotic transcriptional reactivation, but only minimal architectural perturbations. Importantly, the cohesin-independent loops did not show any preferential vulnerability to CBP/p300 inhibition, suggesting the presence of complex compensatory mechanisms beyond CBP/p300 or transcriptional activity. In support of this complexity, recent studies on cohesin-independent looping mechanisms revealed several contributing factors, including the Mediator complex, Polymerase II and transcription itself, LDB1, YY1, and others.^20,53–55,83–87^ In all cases, individual perturbations of any of these factors induced only moderate architectural changes, similar to the ones we uncovered with CBP/p300 inhibition. Therefore, both the maintenance and the establishment of cohesin-independent, enhancer-promoter interactions are likely mediated by multiple active marks and chromatin factors, conferring a complex “molecular memory” resilient to individual perturbations.

In sharp contrast to the extensive and dramatic impact of cohesin depletion on the post-mitotic refolding of TADs and chromatin loops, its effects on the transcriptional reactivation of the ESC program were minimal and comparable to the ones reported in asynchronous cells.^33,39^ Therefore, both the maintenance and the post-mitotic reactivation of transcription are largely independent of cohesin. The efficient transcriptional reactivation in the absence of extruding cohesin could either be driven by the re-formation of cohesin-independent regulatory loops—as discussed above— and/or by the activity of promoters and proximal enhancers that might not require looping.^8,40,41,88^ However, we also identified a limited number of genes, including pluripotency-associated genes (i.e., *Klf4* and *Tbx3*), that showed some degree of transcriptional perturbations upon cohesin depletion during the M-to-G1 transition. These genes had the following characteristics: (i) they were confined within insulated domains (S-S with convergent CTCF motif) and gene-sparse TADs with stronger boundary insulation, (ii) they exhibited higher RAD21 occupancy at promoters, (iii) they were highly connected, and featured many P-S loops. In such contexts, the re-formation of P-E interactions showed higher cohesin-dependency, as supported both by our Micro-C and 4C-Seq data around the example *Klf4* and *Tbx3* genes, contributing to a deficient transcriptional reactivation. Together, our results support that the faithful and efficient transcriptional reactivation of most genes during the M- to-G1 transition shows limited and locus-specific dependency on loop extrusion activity.

The strong enrichment of differentiation-related categories among the few RAD21-dependent genes prompted us to investigate whether the RAD21 depletion during the M-to-G1 transition in the presence of differentiation signals would induce more extensive transcriptional defects. Indeed, the post-mitotic release of synchronized naïve ESCs into EpiLC-inducing conditions in the absence of cohesin impaired the upregulation of many EpiLC-related genes. Although the post-mitotic activation of the majority of genes remained unaffected, the extent of perturbations was substantially greater in the differentiation vs self-renewal conditions and included a large number of important transcription factor genes. This suggests that cohesin-mediated extrusion is more critical for establishing E-P loops and *de novo* transcriptional programs during cell fate transitions rather than maintaining or re-establishing loops in the context of self-renewal. Extending the recently proposed “buffering model”^33^ in self-renewal conditions, persistent chromatin states and mitotic bookmarks could both sustain and establish E-P interactions and transcriptional activity in the absence of cohesin. However, during cell fate transitions, the newly activated enhancers might be more dependent on active extrusion for finding, interacting, and upregulating their target genes. This is in concordance with previous studies reporting cohesin’s important role in inducible gene expression during hematopoietic differentiation.^89–91^ In further support, transient depletion of cohesin in less than one cell cycle significantly impaired the conversion of naïve to formative pluripotency.

In conclusion, by combining acute perturbations of cohesin with cell cycle synchronization and ESC differentiation, our study dissected the degree of functional interdependence between cohesin-mediated chromatin organization and transcriptional activation and revealed unique locus-specific and context-specific vulnerabilities. Importantly, our findings support the presence of a complex “molecular memory” that persists through cell division -during self-renewal conditions- and enables successful re-formation of enhancer-promoter interactions in the absence of cohesin or CBP/p300 activity. In the absence of such “memory”, cohesin extrusion becomes more critical for the establishment of new chromatin interactions and transcriptional programs at the onset of cell fate transitions.

## Methods

### Cell line construction

The *Rad21*-dTAG cell line was derived from Bruce-4 C57BL/6J mouse embryonic stem cells (ESCs; MilliporeSigma, CMTI-2), as previously described.^48^ The KMG *Rad21*-dTAG cell line has a 24×MS2 cassette knocked into the *Klf4* locus and has constitutive expression of MCP-mNeonGreen, as previously described.^48^

### Cell culture

ESCs were cultured at 37°C on surfaces pre-coated with 0.2% w/v gelatin (Sigma-Aldrich, G1890). They were maintained in standard serum/2i/LIF conditions using DMEM (Gibco, 10829018) supplemented with 15% fetal bovine serum (R&D Systems, S10250), 100 U/mL penicillin-streptomycin (Gibco, 15140163), 1X non-essential amino acid (Gibco, 11140076), 1X GlutaMAX supplement (Gibco, 35050079), 55 µM β-mercaptoethanol (Gibco, 92185023), 128 µg/mL LIF (In-house), 1 μM MEK1/2 inhibitor (PD0325901) (Tocris, 4192) and 3 μM GSK3 inhibitor (CHIR99021) (Tocris, 4423).

ESCs were converted to epiblast-like cells (EpiLCs) at 37°C on surfaces coated with 0.2% w/v gelatin (Sigma-Aldrich, G1890) for 3 hours differentiation experiments or with 16.7 μL/mL fibronectin (MilliporeSigma, FC010) for 22 hours differentiation experiments. The differentiation media consisted of a 1:1 mixture of DMEM/F12 (Gibco, 11320082) and Neurobasal media (Gibco, 21103049) supplemented with 1:200 N-2 supplement (Gibco, 17502048), 1:100 B-27 supplement (Gibco, 12587010), 100 U/mL penicillin-streptomycin, 0.375X GlutaMAX™ supplement, 55 µM β-mercaptoethanol, 12.5 ng/mL FGF2 (Gibco, PHG0360), 20 ng/mL Activin A (Gibco, AF-120-14E), and 1% knockout serum replacement (Gibco, 10828010).

### Mitotic arrest and release into G1 using two inhibitors

*Rad21*-dTAG cells were plated as single cells at a density of 40,000 cells/cm^2^. After 24 hours, the cells were treated with 9 µM CDK1 inhibitor (Ro-3306; Selleckchem, S7747) for 16 hours to synchronize them in the G2 phase. Following treatment, the cells were washed with PBS containing 100 ng/mL nocodazole (Sigma-Aldrich, M1404) and incubated in ESC media supplemented with 100 ng/mL nocodazole and 500 nM dTAG-13 (dTAG; Tocris, 6605) for 2 hours to arrest the cells in the M phase and induce RAD21 degradation. 0.05% v/v DMSO (Sigma-Aldrich, D8418) was used as a control. Mitotic cells, distinguished by their rounded morphology and increased susceptibility to mechanical forces, were collected via mitotic shake-off. The collected mitotic cells were washed three times with PBS and subsequently released into the G1 phase for 3 hours, which is approximately the duration of G1 in ESCs. The cells were maintained in either ESC or EpiLC media containing dTAG or DMSO during this period. For CBP/p300 inhibition experiments, 20 µM A485 (Tocris, 6387) was added at the final hour of nocodazole treatment and maintained throughout the experiment, similarly to dTAG. 0.04% v/v DMSO was used as a control.

### Validation of mitotic arrest and release

To validate the purity of mitotic cells, cells resuspended in PBS were placed onto glass surfaces pre-coated with poly-L-lysine (MilliporeSigma, P8920). The cells were centrifuged at 300 × g for 2 min to facilitate attachment to the glass surface. The attached cells were fixed with 2% w/v formaldehyde (Thermo Scientific, 28908) for 10 min, followed by washing with PBS. The cells were permeabilized with 0.5% Triton X-100 (MilliporeSigma, X100) in PBS for 5 min and washed twice with PBS for 3 min each. Blocking was performed with a buffer containing 1% w/v BSA (Gibco, 15260037) in PBS for 30 min. The cells were then incubated with the primary antibody (anti-H3S10ph; Abcam, ab47297), diluted 1:1000 in blocking buffer, for 1 hour. After incubation, the cells were washed twice with PBS and incubated with the secondary antibody (goat anti-rabbit w/ Alexa Fluor 488; Invitrogen, A-11034), diluted 1:500 in blocking buffer, for 45 min. Finally, the cells were washed twice with PBS, stained with 0.3 μg/mL DAPI (Thermo Scientific, 62248) for 3 min, and kept in PBS for imaging. Mitotic purity was calculated as the ratio of H3S10ph-positive foci colocalized with DAPI to the total DAPI-positive foci. Samples with > 90% mitotic purity were used for downstream experiments.

For cell cycle analysis using fluorescence-activated cell sorting (FACS) analysis, 100,000– 500,000 cells were resuspended in 100 μL PBS and fixed by adding 600 μL of 100% ethanol while vortexing the tube at medium speed. The cells were incubated on ice for 10 min, centrifuged at 500 × g for 5 min at 4°C, and the pellet was resuspended in PBS containing 0.3 μg/mL DAPI. The suspension was filtered through 35 μm cell strainer (VWR, 352235) into FACS tubes. Analysis was performed using BD FACS Canto II analyzer and FlowJo v10.10 software. Samples with >80% mitotic release into G1 were used for downstream experiments.

### Western blot

Cells were resuspended in 1X Laemmli butter and sonicated for 5 cycles (30 s on, 30 s off) using the Bioruptor Pico (Diagenode). The samples were then boiled at 95°C for 5 mins. Prepared samples were used for Western Blot analysis with the following antibodies: HA Tag (Invitrogen, 26183), RAD21 (Abcam, ab992), OTX2 (Abcam, ab21990), α-Tubulin (MilliporeSigma, T6199), H3 (Cell Signaling Technology, 9715), H3K72ac (Abcam, ab4927), and H3S10ph (Abcam, 47297).

### RNA-seq

Total RNA was extracted from 500,000 cells per sample using RNeasy Mini Kit (Qiagen, 74104), following the manufacturer’s instructions. For M-to-G1 experiments, RNA-seq libraries were generated by Weill Cornell Medicine Genomics Resources Core Facility using NEBNext Ultra II Directional RNA library kit (New England Biolabs, E7760) and TruSeq Stranded mRNA library kit (Illumina, 20020595). Libraries were sequenced on Illumina NovaSeq 6000 in PE100 mode. For asynchronous experiments, RNA-seq libraries were generated by Novogene using their mRNA library preparation service with poly A enrichment. Libraries were sequenced on Illumina NovoSeq X Plus in PE150 mode.

### ChIP-exo

10,000,000 Bruce-4 cells per replicate were fixed with 1% w/v formaldehyde for 10 min at room temperature (RT) on a platform shaker, followed by quenching with 125 mM glycine for 5 minutes under the same conditions. Crosslinked cells were centrifuged at 500 × g for 5 min at 4°C and were washed twice with ice-cold PBS. After the final wash and centrifuge, the pellet was snap-frozen before the extraction. ChIP-exo libraries were generated by Cornell Biotechnology Resource Center EpiGenomics Core using the ChIP-exo v6.1^93^ protocol. Libraries were sequenced with NextSeq 500 in PE75 mode.

## 4C-seq

For each sample, 2,000,000 cells were crosslinked with 1% w/v formaldehyde for 10 min at RT on a platform shaker, followed by quenching with 125 mM glycine for 5 min under the same condition. The cell pellets were washed twice in PBS and resuspended in 300 μL of lysis buffer (10 mM Tris-HCl pH 8.0, 10 mM NaCl, 0.2% IGEPAL CA-630 (MilliporeSigma, I8896), and 1X cOmplete EDTA-free protease inhibitor cocktail (Roche, 4693132001)) and incubated on ice for 20 min. Following centrifugation at 500 × g for 5 min at 4°C, the pellet was resuspended in 50 μL of 0.5% SDS and incubated at 65°C for 10 min on a shaker at 1,250 rpm. SDS was quenched by adding 150 μL of ddH2O and 25 μL of 10% v/v Triton X-100 (MilliporeSigma, X100) at 37°C for 15 min on a shaker at 1,250 rpm.

To initiate the first digestion, 25 μL of NEBuffer DpnII (New England Biolabs, B0543) and 10 µL of DpnII enzyme (New England Biolabs, R0543) were added, and the samples were incubated overnight at 37°C on a shaker at 750 rpm. The following morning, an additional 2 µL of DpnII was added, and the samples were incubated for 2 hours. Digestion efficiency was confirmed using gel electrophoresis. The enzyme was then inactivated by incubating at 65°C for 20 min. Once the samples cooled down, 645 µL of ddH2O, 120 µL of 10X T4 DNA ligase reaction buffer (New England Biolabs, B0202), 120 µL of 10% v/v Triton X-100, 12 µL of 10 mg/mL BSA (New England Biolabs, B9200), 1 µL of 2000 U/µL T4 DNA ligase (New England Biolabs, M0202), and 60 µL of 10 mM ATP (New England Biolabs, P0756) were added to the samples. The ligation reaction was incubated overnight at 16°C in a water bath in a cold room. The following morning, 0.5 µL of 2000 U/µL T4 DNA ligase and 30 µL of 10 mM ATP were added, and the samples were incubated at RT with rotation until the ligation efficiency was verified using gel electrophoresis. After confirming successful ligation, 60 µL of 20 mg/mL Proteinase K (Fisher Scientific, BP1700) and 120 µL of 10% v/v SDS were added, and the samples were incubated at 65°C for 3 min. Then, 130 µL of 5 M NaCl was added, and the samples were incubated overnight at 65°C on a shaker at 5,000 rpm for reverse crosslinking. The following morning, 10 µL of 10 mg/mL RNase A (MilliporeSigma, R6513) was added, and the samples were incubated at 37°C for 1 hour. DNA was purified using phenol-chloroform extraction and ethanol precipitation, and the pellet was dissolved in 100 µL of 10 mM Tris-HCl pH 8.0.

The second digestion was performed by adding 20 µL of 10X Buffer B (Thermo Scientific, ER0211), 10 µL of 10 U/µL Csp6I, and 80 µL of ddH2O, and incubating overnight at 37°C on a shaker at 700 rpm. After confirming digestion efficiency the next morning, a second ligation was performed by adding 2,204 µL of ddH2O, 300 µL of 10X T4 DNA ligase buffer, 1 µL of 2000 U/mL T4 DNA ligase, and 300 µL of 10 mM ATP. The reaction was incubated overnight at 16°C in a water bath in a cold room. DNA was purified using phenol-chloroform extraction and ethanol precipitation, resulting in the 4C template.

Primers for 4C libraries were designed around the transcription start site (TSS) of each gene of interest according to previously described criteria.^94^ Libraries were generated from the 4C template using the inverse PCR strategy. Briefly, the first round of PCR was performed in quadruplicate per sample, with 150 ng of 4C template used for each reaction in the quadruplicate, employing Expand Long Template PCR system (Roche, 11681834001). The PCR products were pooled to increase the library complexity. A second round of PCR was performed using the initial PCR library as a template, with overlapping primers to add P5/P7 sequencing primers and sample-specific indexes. SPRIselect beads (Beckman Coulter, B23318) were used for DNA size selection throughout the protocol. The libraries were sequenced on Illumina NovoSeq X Plus in PE100 mode. All the primer sequences used for 4C-seq are provided in **Supplementary Table 1**.

### Micro-C

2,000,000 cells per replicate were collected and centrifuged at 300 × g for 5 min. The pellet was frozen at -80°C for 30 min, thawed at RT, and resuspended in 1 mL PBS. Freshly prepared 10 µL of 0.3 M DSG (Sigma-Aldrich, 80424) in DMSO was added, and the samples were incubated with rotation at RT for 10 min. Subsequently, 27 µL of 37% formaldehyde (Sigma-Aldrich, 252549) was added, and the samples were incubated at RT for an additional 10 min. After centrifugation at 3,000 × g for 5 min at 4°C, the pellets were washed with 200 µL of 1X wash buffer from Dovetail Micro-C kit (Cantata Bio, 210006). The pellet was resuspended in 200 µL of MB1 buffer (50 mM NaCl, 10 mM Tris-HCl pH7.5, 5 mM MgCl_2_, 1 mM CaCl_2_, 0.2% v/v NP-40 alternative (MilliporeSigma, 492016), 1 X cOmplete EDTA-free protease inhibitor cocktail in ddH2O) and incubated on ice for 20 min. The samples were centrifuged at 1,750 × g for 5 min at 4°C, and the pellets were washed with 200 µL of MB1 buffer. The nuclei pellets were then resuspended in 200 µL of MB1 buffer, and 28 U of MNase (Worthinigton Biochemical Corporation, LS004798) was added. After brief vortexing, the samples were incubated at 37°C for 20 min on a shaker at 1,000 rpm. The amount of MNase to use was determined based on a separate titration experiment. The samples were transferred onto ice and 1.6 µL of 500 mM EGTA was added to stop the digestion. The reaction was further incubated at 65°C for 10 min for complete inactivation of MNase. The samples were centrifuged at 170 × g for 5 min at 4°C, and the pellets were washed with 1 mL MB2 (50 mM NaCl, 10 mM Tris-HCl pH 7.6, 10 mM MgCl2 in ddH2O). The pellets were resuspended in 50 µL of 1X Nuclease Digestion Buffer from Dovetail Micro-C kit. To lyse the nuclei, 3 µL of 20% v/v SDS was added, and the samples were incubated at 22°C for 15 min on a shaker at 1,250 rpm. The rest of the steps in Micro-C library preparation, including lysate quality control, proximity ligation, library preparation, ligation capture, and library amplification, were performed following Dovetail Micro-C kit user guide v1.2. The libraries were sequenced on Illumina NovoSeq X Plus in PE100 mode.

### Synchronizing cells for single-gene imaging

KMG *Rad21*-dTAG cells were plated on laminin-coated 8-chamber coverglass as previously described.^48,95^ To arrest cells in mitosis, 200 ng/mL nocodazole (Sigma-Aldrich, M1404) was added to the culture media for 6 hours. During the final 2 hours of synchronization, 400 µM dTAG-13 (Sigma-Aldrich, SML2601) was added to induce RAD21 degradation. 0.04% v/v DMSO was used as a control. Cells were released from mitotic arrest by gently washing the chambers 10 times with media lacking nocodazole but containing 400 µM dTAG-13 or 0.04% v/v DMSO as a control). Following the washes, cells were maintained in the same media. Single-gene imaging started within 10 min of the final wash.

### Single-gene imaging

Imaging and quantification of nascent *Klf4* transcription were performed as previously described with few modifications.^48^ 4 different regions were imaged per experiment, capturing an xyz volume of 211 × 211 × 2.5–5 µm^3^ in each region with 270 nm z-steps. Camera exposure times were set to 500 ms at each z position, resulting in a 15-20 s acquisition time per volume for each region. Imaging was conducted over a total duration of 2 hours, with images acquired at 90 s intervals.

### Quantification and statistical analysis

Snakemake (v7.32.4) was used to run pipelines for all genomics analyses.^96^ All statistical analyses and figure generation in this manuscript were done using R (v4.4.0). The Wilcoxon rank-sum test was applied to compare distributions between two groups after confirming that the data did not follow a normal distribution. Fisher’s exact test was used to evaluate differences in composition between the two groups. Two-tailed two-sample Kolmogorov-Smirnov test was used to compare distributions in the cumulative difference function (CDF) plot. Additionally, K-means clustering was used to identify groups of genes with similar expression patterns across the dataset. Quality checks for all genomics assays are provided in **Supplementary Table 1**.

### RNA-seq analysis

FASTQ files were aligned to the mouse genome (mm10) using Star aligner (v2.7.11a)^97^ with the default setting for paired-end reads. Samtools (v1.20)^98^ was used to process the aligned data, including transforming file formats, filtering low-quality reads, and sorting paired reads. FeatureCounts from subread (v2.0.6)^99^ was used to count fragments that have both ends successfully aligned. Normalization of read counts was done in R using RPM and TPM methods. Genes with RPM > 1 in at least 2 samples were considered expressed. Downstream differential analysis was performed with R package DESeq2 (v1.44.0)^100^ using shrunken log2 fold change of <0.5 and adjusted p-value of <0.05 as cutoffs. Batch correction was done using DESeq2 design parameters. Raw read counts, DESeq2 normalized read counts, and DESeq2 analysis are provided in **Supplementary Table 3**.

### ChIP-exo analysis

Adapters were trimmed off from paired-end reads using bbduk.sh (v2.4.5)^101^ with the following option: ktrim=r ref=adapters rcomp=t tpe=t tbo=t hdist=1 mink=11. Filtered reads were aligned to the mouse genome (mm10) using bowtie2 (v2.4.5)^102^ with the options --local --sensitive-local --no-unal. PCR duplicates were removed using MarkDuplicates module from picard (v2.27.4)^103^ with the following option: --VALIDATION_STRINGENCY=LENIENT. Bedtools (v2.30.0)^104^ was used to remove reads overlapping with blacklist region^105^ and convert data to ‘bedgraph’ format. BedGraphToBigWig module from kentUtil^106^ was used to convert data to ‘bigWig’ format. Peaks were called with macs2 (v2.2.7.1) callpeak^107^ with ‘--call-summits’ option.

### 4C-seq analysis

Adapters were trimmed from paired-end reads using cutadapt (v4.8)^108^ with the following options: -j 0 -e 0.15 -m 40. Filtered reads were aligned to the mouse genome (mm10) and normalized using Pipe4C pipeline.^94^ Reads were depth normalized to 1 million cis reads, excluding the top 3 fragments with the highest reads aligned. Smoothening was performed using only non-blind fragments, with sliding bin normalization. Bin sizes of 5 kb were used for *Klf*4, and 10 kb *Tbx3, Myc*, and *Fosl2*, with a bin shift size of 1/20 of the bin size. Visualization of the 4C-seq data was done in R using ggplot2.^109^

### Micro-C analysis

Paired-end reads were aligned to the mouse genome (mm10) using BWA-MEM algorithm from bwa (v0.7.17)^110^ with the following options: -5SP -T0. Parse module of pairtools (v1.1.0)^111^ was used to filter high-quality valid read pairs with the with the following options: --min-mapq 40 --walks-policy 5unique --max-inter-align-gap 30. Dedup module of pairtools (v1.1.0) was used to remove PCR duplicates, followed by split module to generate ‘pairs’ file. Juicer (v1.22.01)^112^ was then used to generate a contact matrix in ‘hic’ format from ‘pairs’ file. For downstream analysis, ‘hic’ format was converted to ‘mcool’ format using HicConvertFormat of HiCExplorer (v3.0.1).^113^ We assessed the reproducibility of the Micro-C data across replicates using GenomeDISCO (v1.0.0)^114^, which computes concordance scores using smoothed contact maps. We calculated concordance scores between replicates as well as across different treatment conditions (DMSO, dTAG, and A495) at a resolution of 10 kb to check the reproducibility of Micro-C. The summary of Micro-C interactions can be found in **Supplementary Table 4**.

Contact probability as a function of genomic separation was calculated using Cooltools (v0.7.0)^115^ in Python, with KR-normalized contact matrices at 1 kb resolution. Cscoretools (v1.1) was used to call A/B compartments at 100 kb resolution, applying a minimum distance threshold of 1 Mb. Saddle plots were plotted using R package GENOVA (v1.0.1)^116^ at 100 kb resolution. The corner score of AA, BB, and AB was calculated based on the signal intensity within a box at 10% of the total saddle plot, positioned at each respective corner. TADs were called based on insulation scores calculated using cooltools (v0.7.0)^115^ in Python, with KR-normalized contact matrices at a resolution of 25 kb and a window size of 125 kb. The TAD boundaries are provided in **Supplementary Table 4**.

To identify loops, we used Chromosight (v.1.6.3)^51^, a computer-vision-based loop-calling algorithm. Chromosight takes a KR-normalized contract matrix at a specific resolution as input and calculates a loop score which is the Pearson correlation coefficient for each pixel of the matrix against a predefined kernel representing a loop structure. The loops were filtered based on a loop score of > 0.3 and a q-value of < 10^-5^. Previous studies have demonstrated that Chromosight outperforms HICCUP algorithm in JUICER^112^ and ‘Call-dots’ function in Cooltools in terms of sensitivity and specificity.^33^ For each pooled Micro-C data per condition, loops were called using ‘detect’ function in Chromosight at resolutions of 25 kb, 10 kb, and 5 kb, with loop-calling options detailed in **Supplementary Table 4**. Loops detected at different resolutions were combined, prioritizing those identified at finer resolutions and discarding overlapping loops from coarser resolutions. To compare loops across conditions, a union set of loops was created by merging loops identified in different conditions using the same strategy. The loop scores at each condition were calculated using ‘detect’ function in Chromosight. A score difference cutoff of 0.2 was used to classify loops as differing across conditions. All loops and their score used in the analysis are provided in **Supplementary Table 4**.

### Virtual 4C (v4C)

Valid read pairs from Micro-C were used to generate virtual 4C profiles. The genome was binned into either 5 kb or 10 kb intervals, and the bin containing the viewpoint was selected. The region of interest was defined as ±2 Mb from the viewpoint bin. Within this region, 5 kb or 10 kb sliding windows with a bin shift size of 1/20 of the bin size were created. Secondary mapped read mates overlapping with sliding bins were counted, and the read counts were normalized to the total sequencing depth of the valid read pairs.

### ChIP enrichment analysis and gene ontology analysis

ChIP enrichment analysis of loop anchors was performed using LOLA (v1.34.0)^52^ with genomic regions within loop anchors that were accessible based on publicly available ATAC-seq for mESC (GSM3106257).^61^ Only ESC-specific data from the LOLA reference database was used. In cases where multiple significant hits were returned for the same target, the result with the highest rank was selected as representative. For analysis of all loops, the most extreme loops from the RAD21- dependent (DOWN) group were selected to match the number of loops in the RAD21-dependent (UP/NO) group. For analysis of regulatory loops, all loops from both the RAD21-dependent (DOWN) and RAD21-independent (UP/NO) groups were used. The full list of ChIP enrichment analysis results is provided in **Supplementary Table 5**. Gene ontology analysis was performed using enrichGO function in clusterProfiler (v.4.12.6).^117^ Benjamini-Hochberg correction with a p-value cutoff of 0.05 and a q-value cutoff of 0.2 was used. The full list of gene ontology analysis results is provided in **Supplementary Table 6**.

## Data availability

All genomic datasets generated in this study, including RNA-seq, ChIP-exo, 4C-seq, and Micro-C, have been deposited in the Gene Expression Omnibus (GEO) under accession number GSE000000. Publicly available genomic datasets used in this study are detailed in **Supplementary Table 1**. Quality control metrics for genomic data, DESeq2 results from RNA-seq, TADs and loops called from Micro-C, ChIP enrichment analysis using LOLA, gene ontology, and K-means clustering are also provided in **Supplementary Table 2–7**.

## Acknowledgments

We sincerely thank all members of the Apostolou and Stadtfeld groups for their critical feedback on this study, and we especially appreciate Alexander Polyzos for his guidance on computational analysis. We also thank Gerd A. Blobel and Anders Sejr Hansen for their valuable input and David M. Gilbert for his advice on cell cycle synchronization using two inhibitors. Additionally, we are thankful to the reviewers for their insightful and constructive comments, which significantly improved the manuscript. U.L. was supported by the Korea Foundation for Advanced Studies (KFAS) Overseas PhD Scholarship. This study was supported by the Tri-Institutional Stem Cell Initiative (TriSCI), funded by the Starr Foundation, and by the National Institute of General Medical Sciences (NIGMS) (RM1GM139738 and R01GM138635).

## Author contributions

E.A. conceived, designed, and supervised the whole study and analysis with input from U.L., C.L., and A.P. U.L. performed all experiments and analyses in this study, except for single-gene imaging. A.L.-D. assisted with cell cycle synchronization and EpiLC differentiation experiments. B.P.-W. assisted with initial synchronizations, perturbations, and 4C-seq experiments. Z.N. developed the synchronization assay for single-gene imaging, performed the experiments, and analyzed the resulting imaging data under the supervision of A.P. L.C. and J.L. created and validated the cell lines used in the study, with L.C. also conducting preliminary single-gene imaging experiments. W.W. assisted with 4C-seq analysis with guidance from C.L. U.L. prepared all figures and, together with E.A., wrote the manuscript with input from all authors.

---

**Extended Data Fig. 1.**
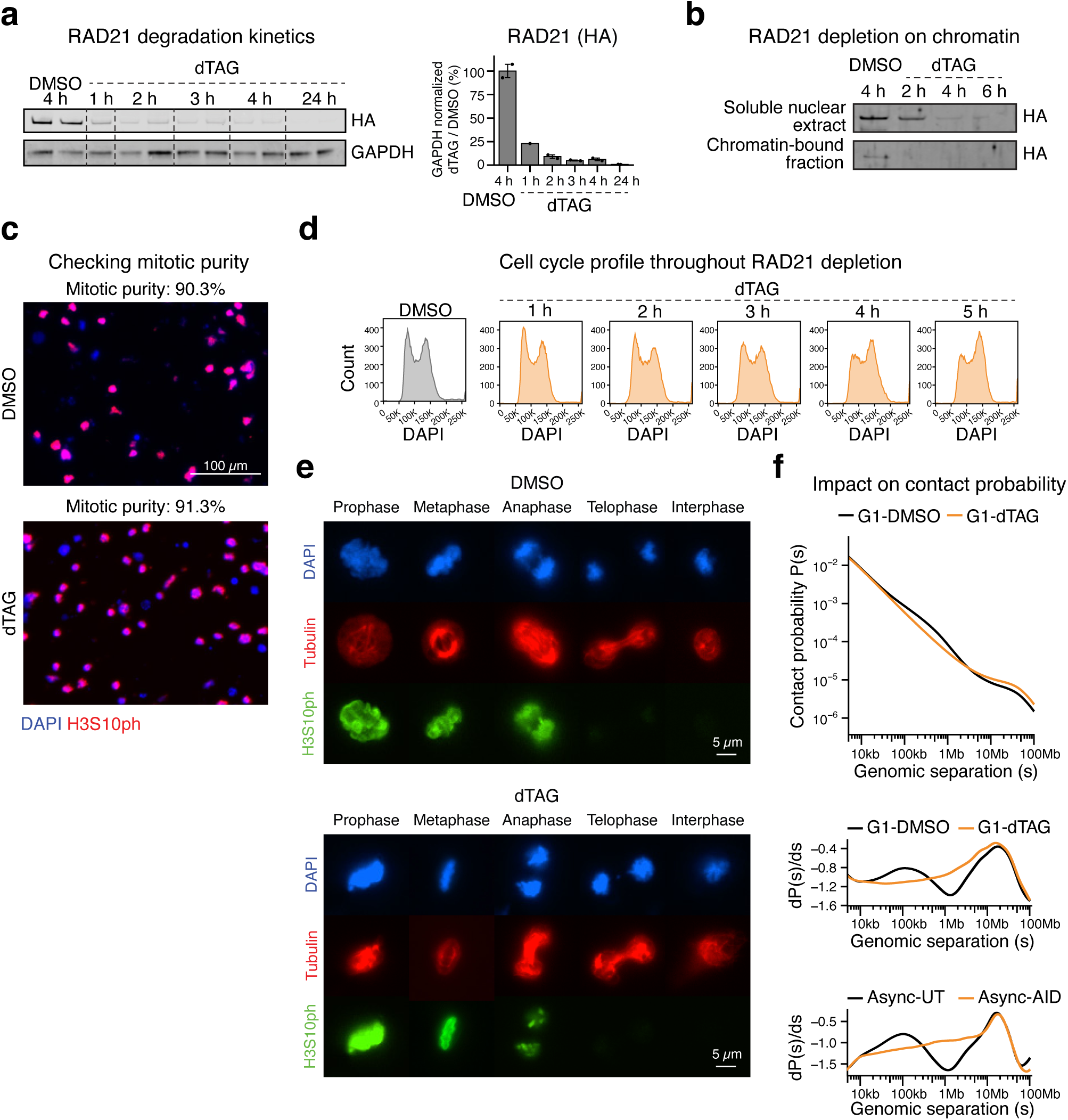
| Related to. Figure 1**. a**, Western blot images showing the time-course depletion of RAD21 (anti-HA) in asynchronous ESCs upon dTAG treatment. **b**, Western blot showing RAD21 (anti-HA) depletion in both the soluble nuclear extracts and the chromatin-bound fraction of ESCs upon dTAG treatment. **c**, Immunofluorescence staining of ESCs, synchronized at mitosis, using DAPI and anti-H3S10ph (a mitotic marker) to evaluate mitotic purity before release. Mitotic purity was calculated as the ratio of H3S10ph-positive foci colocalized with DAPI to the total DAPI-positive foci. **d**, Cell cycle profile analyzed by FACS using DAPI staining in asynchronous ESCs treated with DMSO or dTAG over a time course. **e**, Immunofluorescence staining of ESCs in prophase, metaphase, anaphase, telophase, and interphase using DAPI, anti-Tubulin, and anti-H3S10ph upon DMSO or dTAG treatment. **f**, Top panel: Line plot showing the contact probability (P(s)) as a function of genomic separation (s) around 5 kb to 100 Mb range in G1 Micro-C data upon RAD21 depletion. A bin size 1 kb was used. Bottom panel: Line plot showing the derivative of contact probability (dP(s)/ds) as a function of genomic separation (s) in Micro-C data for G1 (this study) and asynchronous (async) ESCs.^33^

**Extended Data Fig. 2.**
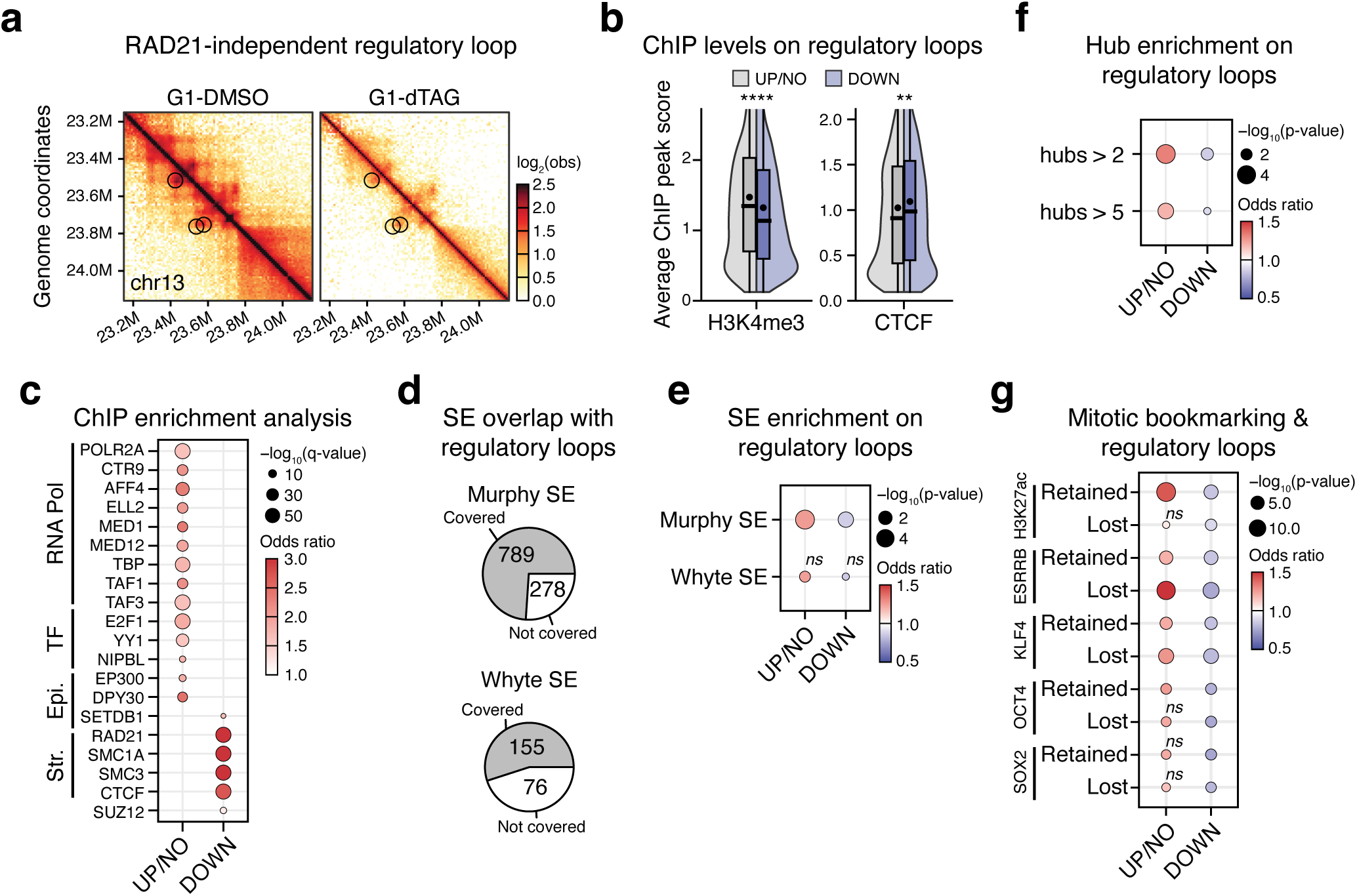
| Related to. Figure 2**. a**, Snapshot of Micro-C contact matrix showing the observed (obs) contact frequency from G1-DMSO and G1-dTAG conditions highlighting RAD21- independent regulatory loops in circles. **b**, Violin and box plot showing the distribution of H3K4me3 and CTCF ChIP-exo peak scores at the anchors of regulatory loops (P-P, P-E, E-E) that are weakened (DOWN) or not (UP/NO) upon RAD21 depletion (as defined in **panel f**). Boxes represent the interquartile range, lines indicate the median, and dots represent the mean. Statistical significance was assessed using the Wilcoxon test (**p < 0.01, ****p < 0.0001). **c**, Dot plot showing top enriched features for the regulatory UP/NO or DOWN loops (as defined in Fig. 2f) based on ChIP enrichment analysis using the LOLA algorithm at the respective loop anchors. The analysis was limited to accessible regions defined by ATAC-seq peaks^61^ within regulatory loop anchors. Only features with significant enrichment in either loop category (q-value < 0.05) are displayed. Dot size represents −log10(q-value), while color represents the odds ratio of enrichment. A complete list of significant results is provided in **Supplementary Table 5**. **d**, Pie chart showing the proportion and numbers of regulatory loops that overlap with super-enhancers (SE).^13,60^ **e**, Dot plot showing the relative enrichment of superenhancers (SE)^13,60^ between RAD21-dependent (DOWN) and independent (UP/NO) regulatory loops (as defined in Fig. 2f). Statistical significance was assessed using Fisher’s exact test (*ns* = not significant). Dot size represents -log_10_(p-value), while color represents the odds ratio of enrichment. **f**, Dot plot showing the relative enrichment of hubs^13^ with more than 2 or 5 connections between RAD21-dependent (DOWN) and independent (UP/NO) regulatory loops (as defined in Fig. 2f). Statistical significance was assessed using Fisher’s exact test. Dot size represents −log10(p-value), while color represents the odds ratio of enrichment. **g**, Dot plot showing the relative enrichment significance of mitotically bookmarked (retained) and non-bookmarked (lost) ChIP-seq peaks for various bookmarking factors (H3K27ac, ESRRB, KLF4, OCT4, and SOX2)^42,64,92^ between RAD21-dependent (DOWN) and independent (UP/NO) regulatory loops (as defined in Fig. 2f). Statistical significance was assessed using Fisher’s exact test (*ns* = not significant). Dot size represents −log_10_(p-value), while color represents the odds ratio of enrichment.

**Extended Data Fig. 3.**
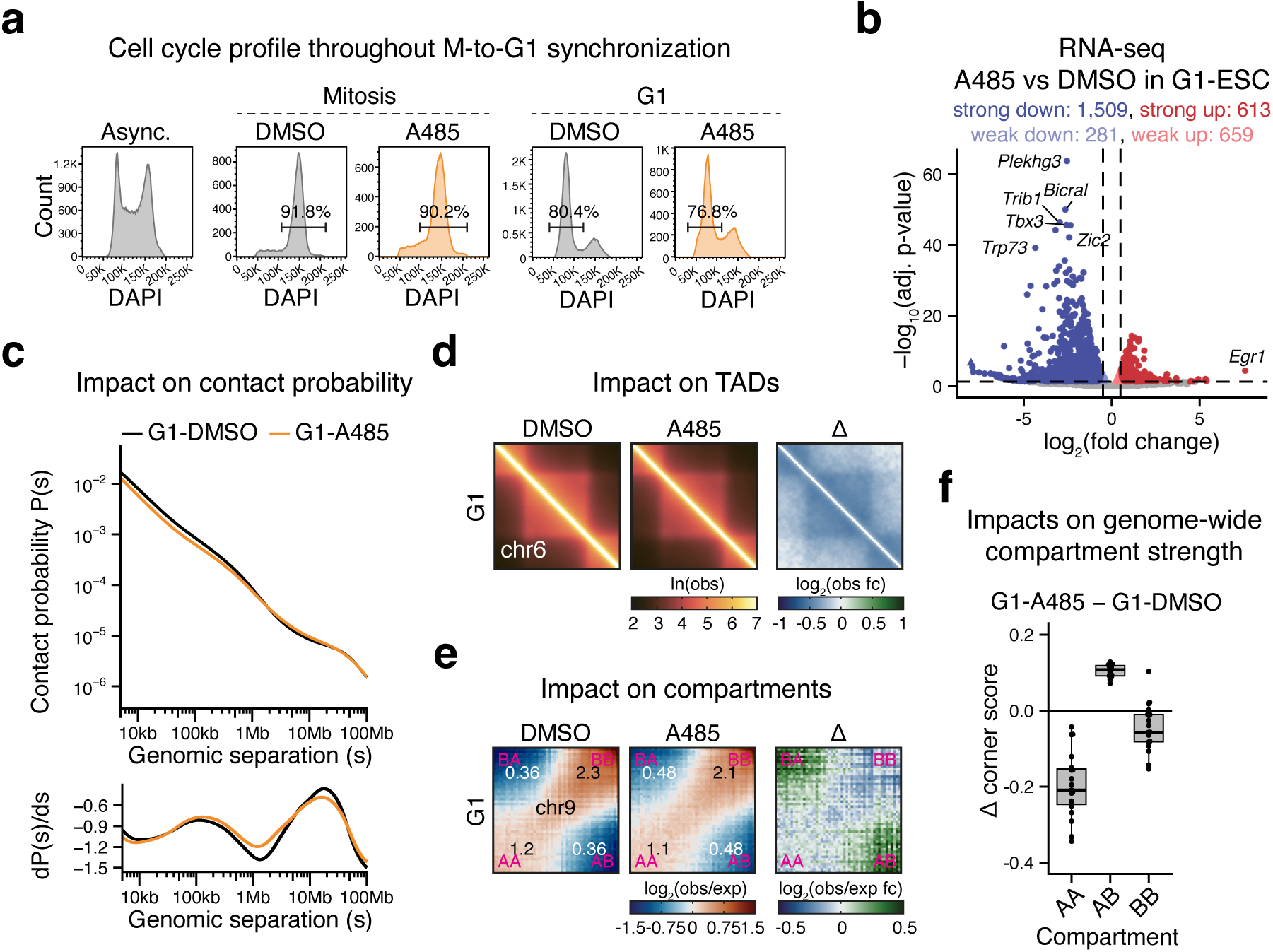
| Transcriptional and architectural changes upon p300/CBP inhibition with A485 treatment. **a**, Cell cycle profile analyzed by FACS using DAPI staining, highlighting a prominent 4N peak (with percentage) in mitosis and a prominent 2N peak (with percentage) in G1, comparing A485-treated ESCs with the DMSO control. **b.** Volcano plot of RNA-seq data highlighting transcriptional changes upon A485 treatment in G1-ESC. Statistical significance was determined using an adjusted p-value cutoff of 0,05, and changes were classified as strong or weak based on an absolute shrunken log_2_ fold change cutoff of 0.5. The analysis was performed using n=2 replicates per each condition. **c**, Line plot showing the contact probability (P(s), *top*) as a function of genomic separation (s) and its derivative (dP(s)/ds, *bottom)* around 5 kb to 100 Mb range in G1 Micro-C data upon CBP/p300 inhibition. A bin size of 1 kb was used. **d**, Left panel: Representative aggregate TAD plots from chr6 in DMSO- and A485-treated G1 ESCs. Color represents natural log (ln) of the observed (obs) contact frequency. Right panel: Differential (Δ = A485 − DMSO) plot comparing A485- treated and control conditions. Color represents log_2_ fold change in observed (obs) contact frequency. **e**, Left panel: Representative saddle plots from chr9 in DMSO- and A485-treated G1 ESCs. The average scores for the 10% area at each corner are displayed. Color represents log_2_ of observed (obs) contact frequency over expected (exp) contact frequency. Right panel: Differential (Δ = A485 − DMSO) plot comparing A485-treated and control conditions. Color represents log_2_ fold change in observed (obs) contact frequency over expected (exp) contact frequency. **f**, Box plot showing the intra- (AA or BB) or inter (AB) compartmental changes across all chromosomes upon CBP/p300 inhibition. The y-axis shows the difference (Δ = A485 − DMSO) of the saddle plot corner scores (as shown in **panel e**). Boxes represent the interquartile range, with lines indicating the median. Each data point corresponds to an individual chromosome.

**Extended Data Fig. 4.**
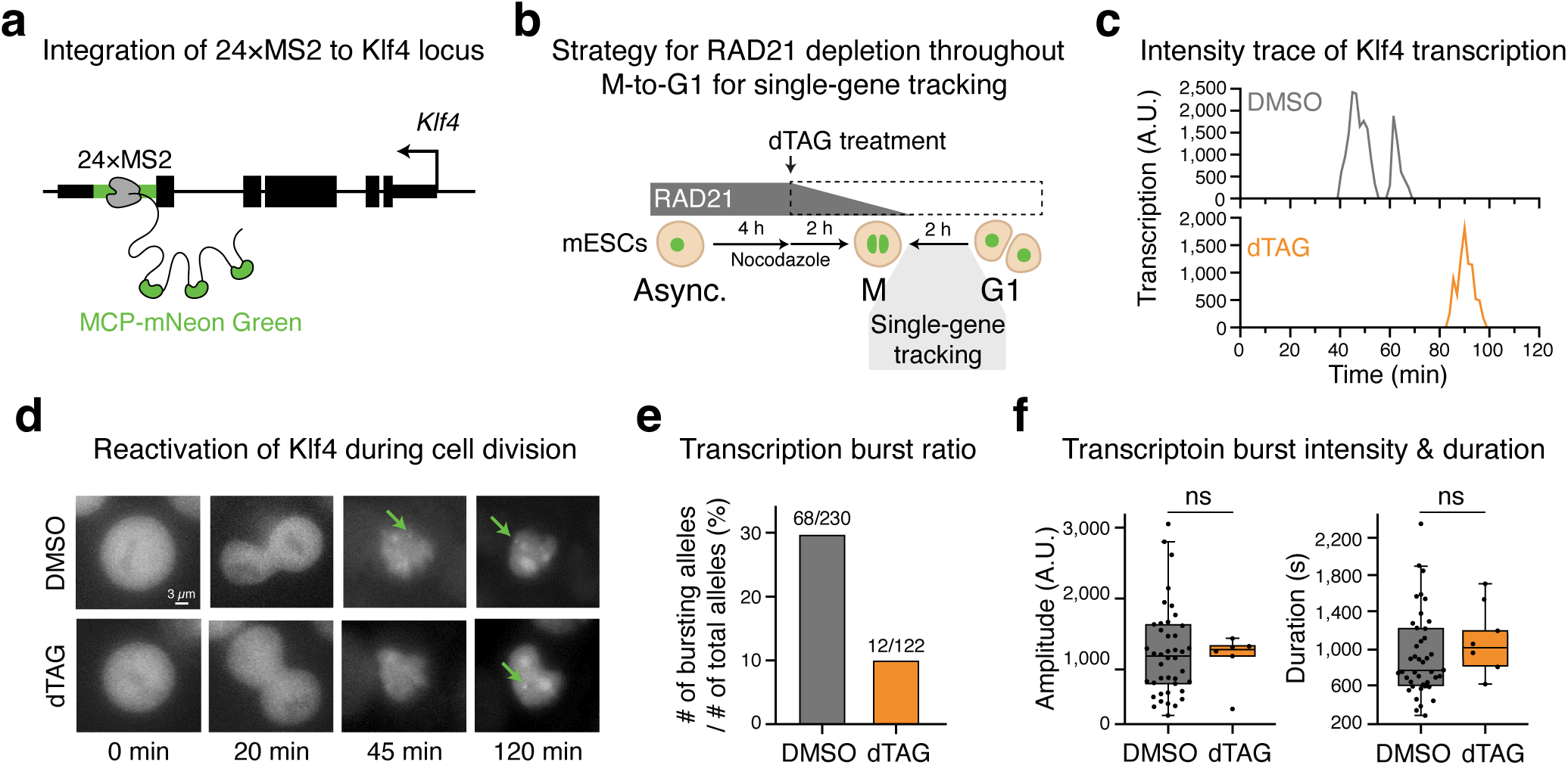
| Single-gene imaging of *Klf4* during mitotic exit upon RAD21 depletion. **a**, Schematic of the integration of 24 × MS2 into the *Klf4* locus of mouse ESCs. MS2 RNA stem-loops are recognized by MCP protein fused to mNeonGreen fluorescent protein for visualization. Black rectangles: exons and 5’/3’-UTR. Green rectangles: 24 × MS2. **b**, Schematic of cell synchronization for single-gene imaging experiments. Cells were synchronized in mitosis by adding 200 ng/mL nocodazole for 6 hours and washed with nocodazole-free media to release from mitosis. 400 µM dTAG was used to deplete RAD21, with 0.4% v/v DMSO used as control. **c**, Live-cell snapshots from single-gene imaging experiments showing reactivation of *Klf4* transcription during cell division. The yellow arrows point to MS2-MCP-tagged *Klf4* nascent transcription sites. **d**, Example intensity traces of *Klf4* transcription sites from DMSO- and dTAG-treated cells. **e**, Ǫuantification of *Klf4* alleles that show transcriptional bursts in the 2-hour imaging period after mitotic exit. Data are driven from two independent experiments. The number of alleles showing bursts out of the total alleles imaged is 68/230 for DMSO and 12/122 for dTAG. **f**, Box plots showing the burst amplitude (left panel) and burst duration (right panel). Boxes represent the interquartile range and lines indicate the median. Each dot represents a single burst. Statistical significance was assessed using the Wilcoxon test (*ns* = not significant).

**Extended Data Fig. 5.**
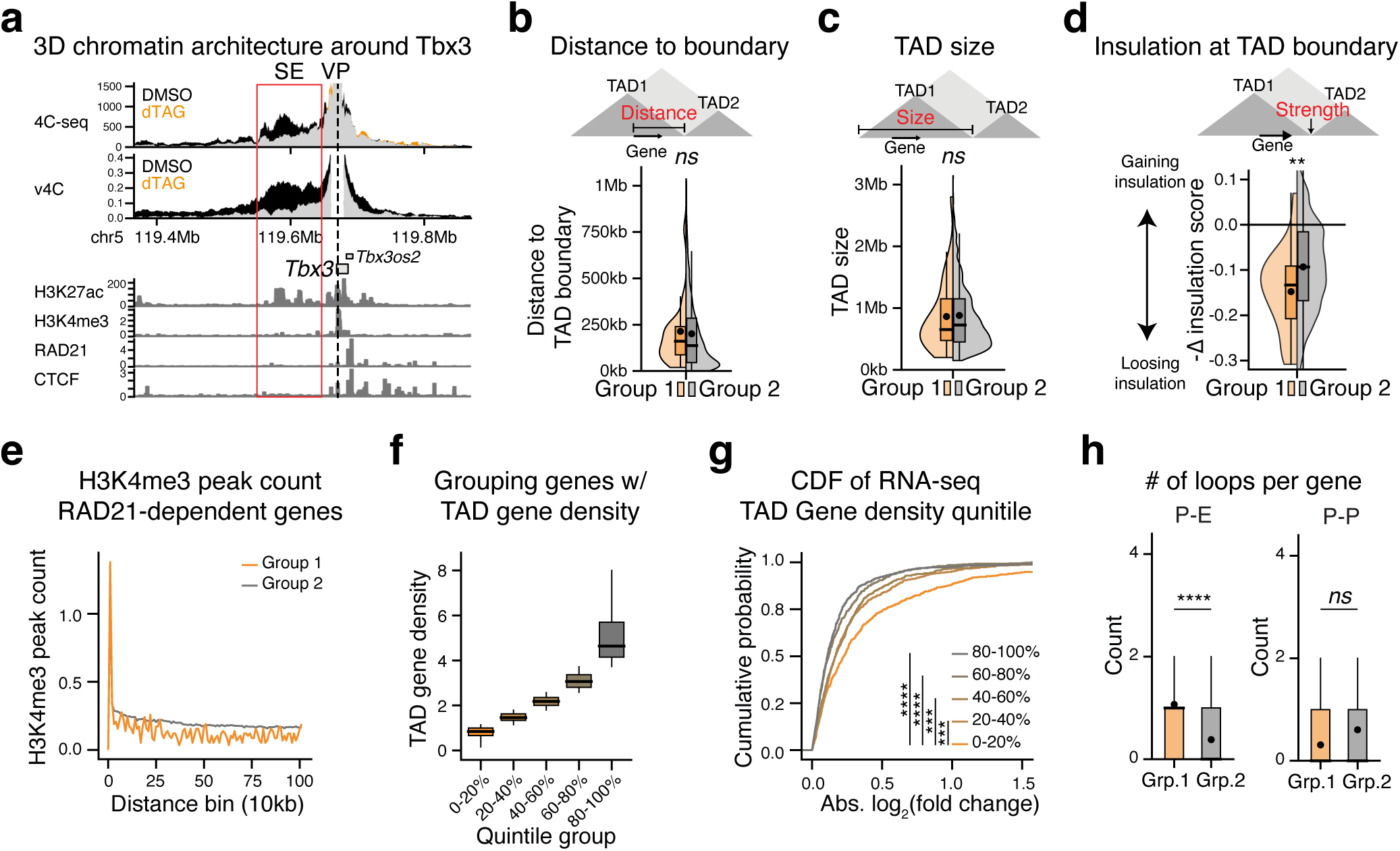
| Related to. Figure 3**. a**, Top panel: Experimental 4C-seq data analysis of ESCs in G1 treated with DMSO or dTAG throughout the M-to-G1 transition, shown as average CPM around the viewpoint (VP, *Tbx3* promoter). Regions with higher 4C-seq signals in DMSO are shown in black, while those with higher signals in dTAG are shown in orange. Middle panel: Virtual 4C calculated from Micro-C of ESCs in G1 with DMSO or dTAG treatment, represented similarly. Bottom panel: ChIP-seq (H3K27ac) and ChIP-exo (H3K4me3, RAD21, and CTCF) tracks. The superenhancer (SE) upstream of the *Tbx3* promoter is highlighted in red. **b**, Box plot and violin plot showing the distance of genes to the closest TAD boundary for genes in Group 1 (orange) or Group 2 (grey). Boxes represent the interquartile range, lines indicate the median, and dots represent the mean. Statistical significance was assessed using the Wilcoxon test (*ns* = not significant). **c**, Box plot and violin plot showing the size of TADs where genes in Group 1 (orange) or Group 2 (grey) genes are located. Boxes represent the interquartile range, lines indicate the median, and dots represent the mean. Statistical significance was assessed using the Wilcoxon test (*ns* = not significant). **d**, Box plot and violin plot showing the negative value of differential (Δ) insulation score (dTAG vs DMSO) of the closest TAD boundary to the genes in Group 1 (orange) or Group 2 (grey). A more negative Δ insulation score indicates a greater loss of TAD boundary insulation upon dTAG treatment. Boxes represent the interquartile range, lines indicate the median, and dots represent the mean. Statistical significance was assessed using the Wilcoxon test (*ns* = not significant). **e**, Line plot showing the average number of H3K4me3 ChIP-exo peaks at various linear distances from the TSS of genes in Group 1 (orange) and Group 2 (grey). The number of peaks was quantified per 10 kb bin. **f**, Box plot showing the gene density distribution across quantile groups, ranked based on TAD gene density (calculated as the number of genes divided by TAD size). **g**, Cumulative distribution function (CDF) plot showing the absolute log_2_ fold change of RNA-seq (dTAG vs DMSO in G1-ESC), measuring the overall transcriptional perturbation upon RAD21 depletion during the M-to-G1 transition. Genes are grouped based on the gene density within TAD (as defined in **panel f**). Statistical significance was assessed using a two-tailed, two-sample Kolmogorov-Smirnov test (***p < 0.001, ****p < 0.0001). **h**, Box plot showing the number of P-E loops and P-P loops per gene in Group 1 (orange) and Group 2 (grey). Boxes represent the interquartile range, lines indicate the median, and dots represent the mean. Statistical significance was assessed using the Wilcoxon test (*ns* = not significant, ****p < 0.0001).

**Extended Data Fig. 6.**
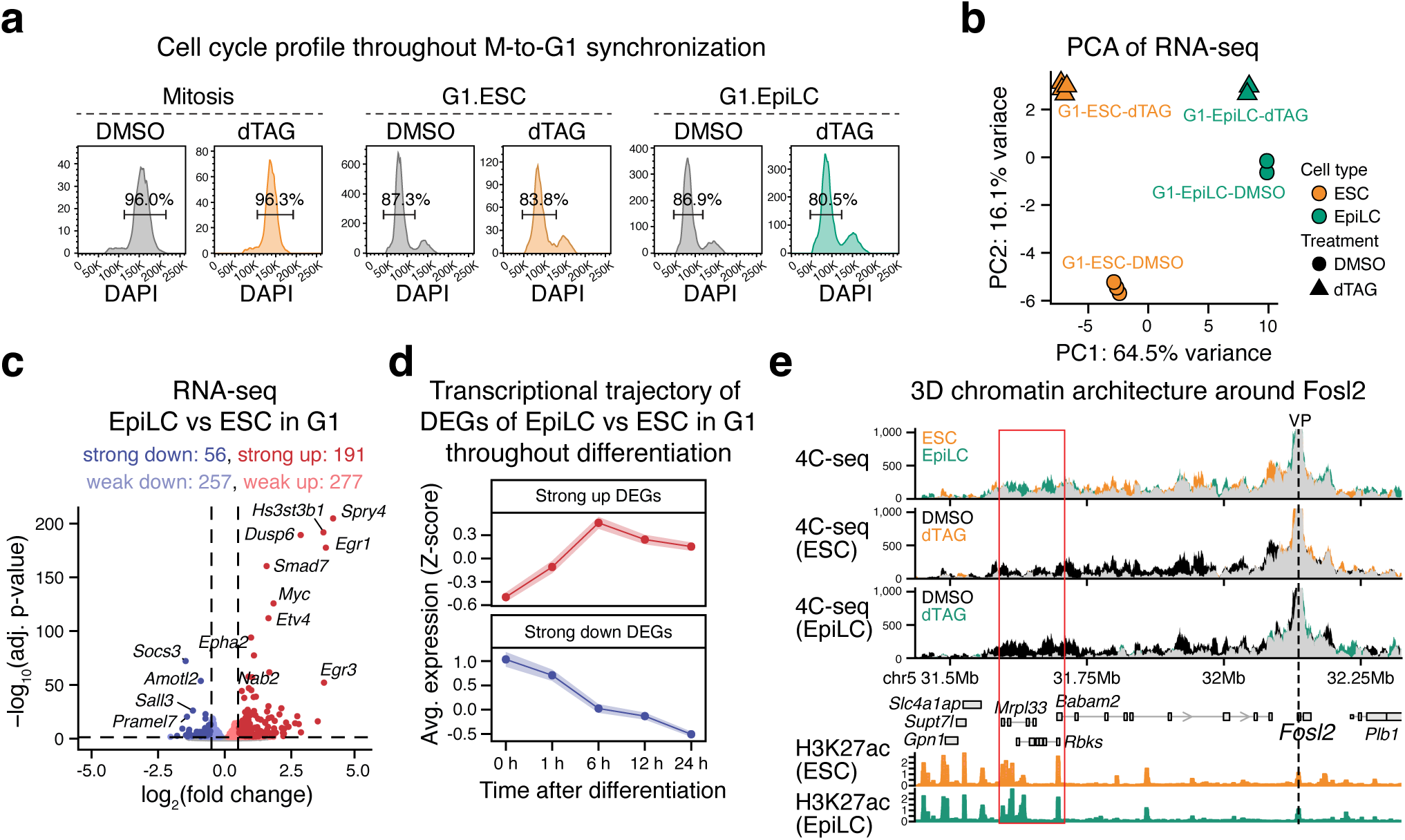
| Transcriptional and architectural changes upon RAD21 depletion in G1 during EpiLC differentiation. a, Cell cycle profile analyzed by FACS using DAPI staining, highlighting a prominent 4N peak (with percentage) in mitosis and a prominent 2N peak (with percentage) in G1, comparing dTAG-treated cells with the DMSO control. **b**, PCA plot of RNA-seq from the G1 population in ESCs and EpiLCs, with or without dTAG treatment. The top 25% of genes with the highest variance were used for plotting. **c**, Volcano plot of RNA-seq data highlighting transcriptional changes upon EpiLC differentiation in G1. Statistical significance was determined using an adjusted p-value cutoff of 0,05, and changes were classified as strong or weak based on an absolute shrunken log_2_ fold change cutoff of 0.5. The analysis was performed using n=3 replicates for DMSO and n=2 replicates for EpiLC. **d**, Line plot showing the transcriptional trajectory of strong DEGs from RNA-seq comparing EpiLC to ESC in G1 (as shown in **panel c**), based on a published dataset.^72^ The average z-score and its standard error (as ribbon) at each timepoint are plotted. **e**, Experimental 4C-seq data analysis comparing interaction patterns around the *Fosl2* promoter viewpoint (VP) between ESC vs EpiLC in DMSO conditions (top panel) or DMSO vs dTAG in ESC conditions (middle panel) or EpiLC conditions (bottom panel). 4C-seq signal is expressed as normalized counts per million (CPM). Regions with higher 4C-seq signals in DMSO are shown in black, while those with higher signals in dTAG are shown in orange. The H3K27ac ChIP-seq tracks for ESC and EpiLC are displayed at the bottom panel. The red box highlights a putative regulatory region >600 kb away from the *Fosl2* promoter which exhibits increased contacts in EpiLC compared to ESC conditions and undergoes EpiLC-specific perturbation upon dTAG treatment.

**Extended Data Fig. 7.**
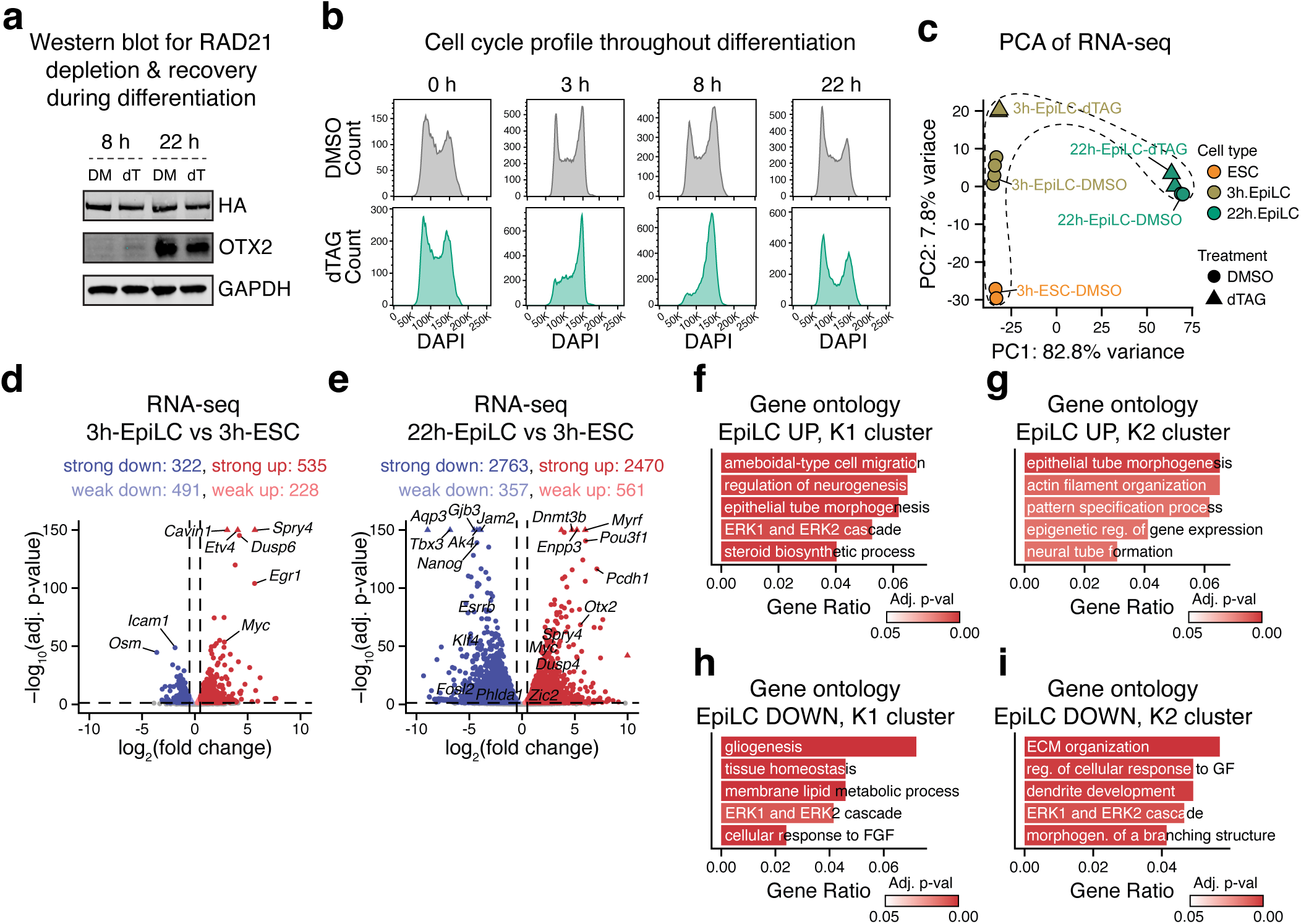
| Transcriptional changes throughout EpiLC differentiation in asynchronous population upon RAD21 depletion and recovery. a, Western blot showing recovery of RAD21 (HA-tagged) during EpiLC differentiation. Also shows OTX2 induction during EpiLC differentiation which is partially perturbed by dTAG treatment. b, Cell cycle profile analyzed by FACS using DAPI staining in asynchronous cells throughout EpiLC differentiation with or without dTAG treatment. c, PCA plot of RNA-seq during EpiLC differentiation with or without dTAG treatment. The top 25% of genes with the highest variance were used for plotting. d, e, Volcano plot of RNA-seq data highlighting transcriptional changes upon EpiLC differentiation at 3 hours (d) and 22 hours (e). Statistical significance was determined using an adjusted p-value cutoff of 0,05, and changes were classified as strong or weak based on an absolute shrunken log_2_ fold change cutoff of 0.5. The analysis was performed using n=2 replicates except for the EpiLC-3h condition which had 4 replicates. f–i, Bar plot showing select biological process significantly enriched in K1 cluster (f), K2 cluster (g) of EpiLC UP genes, and K1 cluster (h) and K2 cluster (i) of EpiLC DOWN genes. Color represents adjusted p-value. An adjusted p-value cutoff of 0.05 and a q-value cutoff of 0.2 were used to determine significance. A complete list of significant results is provided in Supplementary Table 6.

